# Prediction of SUMOylation targets in Drosophila melanogaster

**DOI:** 10.1101/2022.08.19.504577

**Authors:** Yogendra Ramtirtha, M. S. Madhusudhan

**Author notes:** To whom correspondence should be made.

## Abstract

SUMOylation is a post translational modification that involves covalent attachment of SUMO C-terminus to side chain amino group of lysine residues in target proteins. Disruption of the modification has been linked to neurodegenerative diseases and cancer. Recent improvements in mass spectrometry-coupled proteomics experiments have enabled high-throughput identification of SUMOylated lysines in mammalian cells. One such study was Hendriks et al, 2018, wherein the authors identified SUMOylated lysines in human and mouse cells. Information from this study was used as an input to a sequence homology based method to annotate putative SUMOylatable lysines from the proteome of fruit fly *Drosophila melanogaster*. 5283 human and 468 mouse SUMOylated proteins led to the identification of 8539 and 1700 fly homologs and putative SUMOylation sites therein respectively. Clustering analysis was carried out on these annotated sites to obtain three typs of information. First type of information revealed amino acid preferences in the local sequence vicinity of the annotated sites. This exercise confirmed that ψ – K – x – (E/D) where ψ = I/V/L, is the most frequently occurring sequence motif involving SUMOylated lysines.

Second type of information revealed protein families that contain the annotated sites. Results from this exercise reveal that members of thousands of protein families contain annotated SUMOylation sites. Third type of information revealed preferred biological and cellular functions of proteins containing the annotated lysines. This exercise revealed that nucleus and transcription are preferred cellular localization and biological function respectively.

## 1 Introduction

Small Ubiquitin-related MOdifier (SUMO) is a post-translational modifier protein of ∼100 amino acids conserved in all eukaryotes from yeast to humans. SUMO is structurally similar to ubiquitin. Both the proteins belong to the beta-grasp fold of proteins. The SUMO pathway contains SUP (SUMO-specific Proteases), E1, E2 and E3 enzymes. Covalent attachment of SUMO C-terminus to NZ atom of lysine residue in target proteins is known as SUMOylation. SUMOylation has been shown to regulate various biological processes such as nucleo-cytoplasmic translocation of proteins and the activity of transcription factors. Disruption of SUMOylation has been linked to various neuro-degenerative diseases and cancer (a detailed review of SUMOylation can be seen in [1]).

In the last few years, development of mass-spectrometry coupled proteomics experiments has enabled identification of SUMOylation sites in thousands of human proteins (a detailed review can be found in [8]). In addition to standard cellular growth conditions, these experiments also probed the SUMOylation status of proteomes from cells grown under different stress conditions such as heat shock and proteasome inhibitors. Recently, [9] have come up with a list of human and mouse SUMOylated lysines and proteins using mass-spectrometry based proteomics approach.

SUMO-proteomics experiments in the fruit fly *Drosophila melanogaster* have studied the cellular effects of the modification [2–4]. These experiments identified SUMOylated proteins but technical difficulties hindered the identification of modified lysines in these proteins. Knowledge of SUMOylated lysines is important for understanding biological implications of the modification.

There are 2 computational approaches to predict putative SUMOylation sites. First method involves using currently available SUMOylation site prediction tools such as GPS-SUMO [5, 6] and JASSA [7]. These tools use protein sequence as input and scan local sequence environment around lysine residues to make predictions. However, these tools miss information about known SUMOylated lysines from homologous proteins. Second method to predict putative SUMOylation sites is presented in this study. The proposed method begins by identifying fruit fly homologs of known SUMOylated proteins from other organisms. Homology search is based on protein sequence alignments. Putative SUMOylation sites are annotated based on quality of sequence alignments and local sequence environment around lysine residues.

In this study, human and mouse SUMOylated proteins were used to identify orthologous proteins from the proteome of fruit fly *Drosophila melanogaster*. Homology search was carried out using the sequence alignment tool PSIBLAST (Position Specific Iterative Basic Local Alignment Search Tool). In addition, sequence patterns involving SUMOylation sites were studied. Two kinds of information were obtained from sequence pattern analysis. First kind of information was obtained by analyzing local sequence around SUMOylated lysines to detect new motifs. Second kind of information was extracted by clustering target proteins according to their families and identifying lysines conserved in every family. Apart from sequence patterns, Gene Ontology analysis was also carried out to study preferred cellular compartments and biological processes of the modified proteins.

## 2 Materials and Methods

### 2.1 Overview of the method used to identify fly orthologs

The objective of this work was to identify fly homologs of human and mouse SUMOylated proteins. For this reason, human and mouse proteins were queried against a reference database containing the fruit fly proteome and UniRef90 proteins using PSIBLAST. All the details concerning the list of human proteins, reference database and PSIBLAST parameters will be discussed in the following subsections. Alignments involving fly-human and fly–mouse protein pairs were extracted from PSIBLAST results. All the pair-wise alignments were scanned to check whether SUMOylated lysines from human proteins were aligned to a lysine residue from the corresponding fly protein. If this was the case, then 15-mers centered on the lysines of interest were extracted. The 15-mers were extracted by including 7 residues upstream and downstream with respect to lysine of interest. There were 2 lists of 15-mers per organism. The first list was extracted from the FASTA protein sequences (referred to as FASTA 15-mers). And the other list was extracted from the aligned region of the protein sequence (referred to as Aligned 15-mers, which may contain gaps from alignments). The workflow for obtaining 15-mers is summarized below (Figure-1).

**Figure 1:**
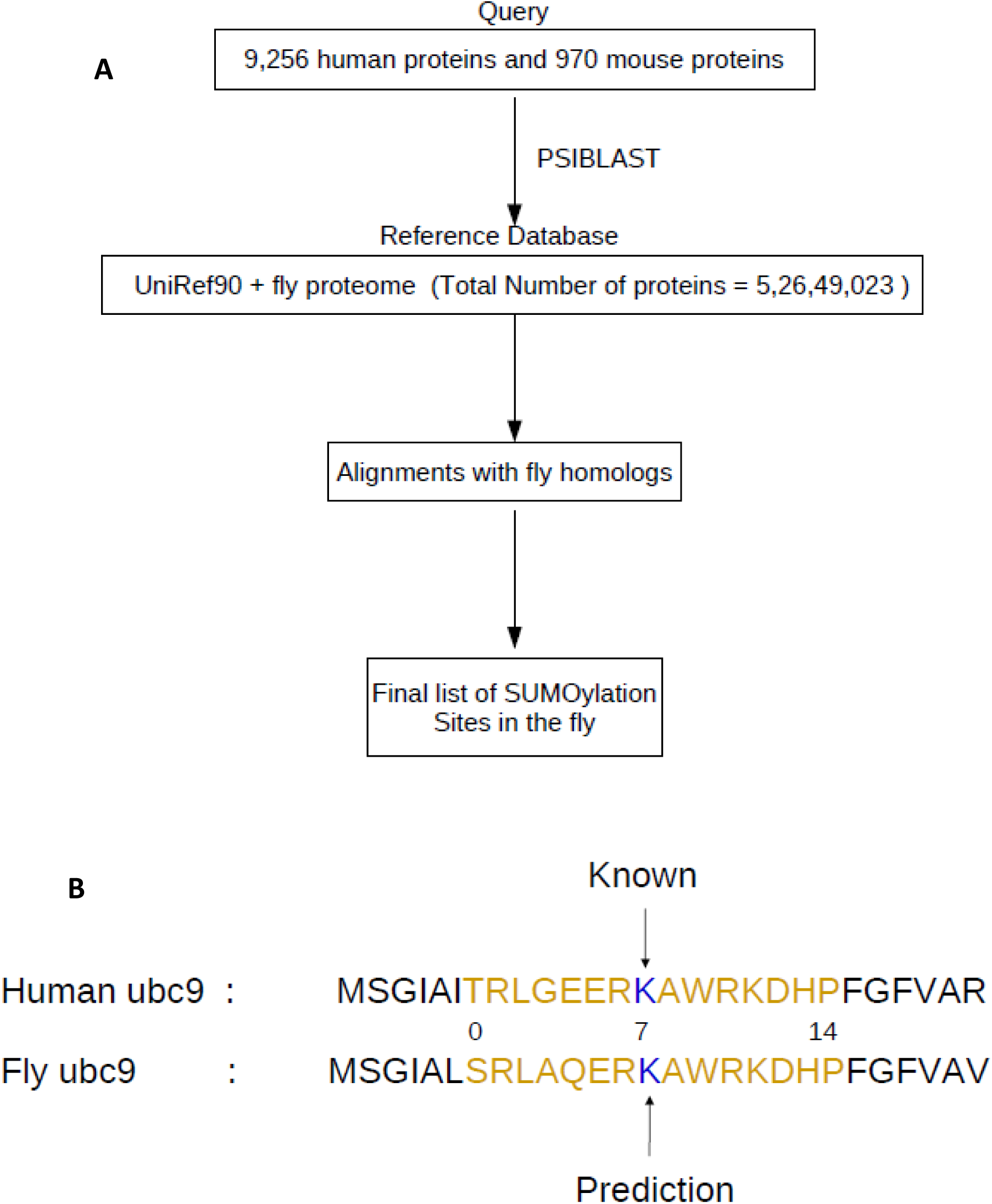
A: Overview of methods used to identify fly orthologs and 15-mers centered on aligned SUMOylated lysines using human and mouse proteins with known SUMOylated lysines. The homology search was carried out using PSIBLAST. B: An example alignment between human and fly homologs of the SUMO E2 conjugating enzyme (ubc9) and the conserved lysines as well as 15-mers therein. The alignment shown here has an E-value of 1e-78.

### 2.2 Reference database and query proteins

For this study, the list of human and mouse SUMOylated proteins was obtained from supplementary data of [9] (human data file named -

41467_2018_4957_MOESM6_ESM.xlsx and mouse data file named -

41467_2018_4957_MOESM8_ESM.xlsx) . Details of query proteins used in this study are summarized below (Table-1).

**Table-1:** Details of query proteins used in this study

| Query type | Number of proteins | Source Organism | Reference |
| --- | --- | --- | --- |
| 1 | 9256 | Human | [9] |
| 3 | 964 | Mouse | [9] |

The FASTA sequences of all the query proteins and reference databases were downloaded from the UniProt database (https://www.uniprot.org/) [10].

The reference database (5,26,49,023 proteins) used in this study was created by combining UniRef90 database (5,26,29,880 proteins) and fruit fly proteome (19,143 proteins, Tax ID – 7227). In order to remove duplicates in the reference database, protein sequences in the UniRef90 database that came from the fruit fly proteome (TaxID – 7227) and having the term “Drosophila melanogaster” in their header were removed.

Homologs of query proteins were searched in the reference proteome using PSIBLAST version 2.7.1+ [11, 12]. For every query protein, 2 kinds of PSIBLAST jobs were carried out. The parameters used for both kinds of jobs are summarized below (Table-2).

**Table-2:**
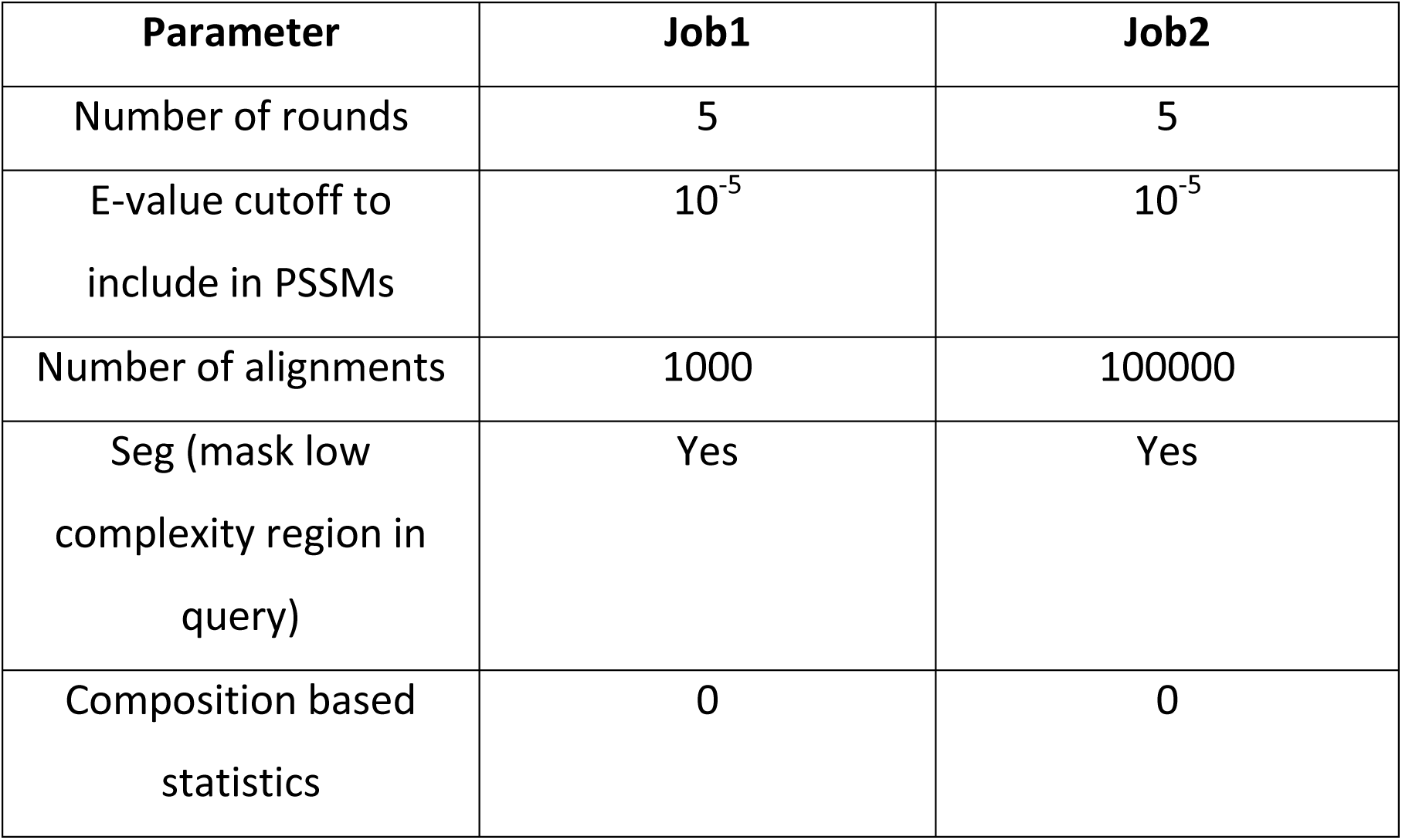
Parameters of both PSIBLAST jobs

For every query protein, 2 kinds of PSIBLAST jobs were carried out. The PSSMs (Position Specific Scoring Matrices) generated after both the jobs differ because of the difference in the number of alignments used namely – 1,000 and 100,000 respectively. The job with 1,000 alignments was carried out to detect close homologs, whereas the job with 1,00,000 alignments was carried out to detect distant homologs. Pair-wise alignments in different rounds of PSIBLAST use different PSSMs. Thus, E-values from different rounds for the same protein pair cannot be compared with each other because PSIBLAST takes PSSMs into account while calculating E-values. Hence, E-values for all pair-wise alignments between human – fly and mouse – fly proteins were re-calculated using BLOSUM62 substitution matrix and the E-value equation (Equation 1).

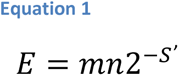

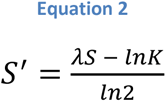

Where E = E-value, S = raw score calculated using BLOSUM62 substitution matrix, S’ = bit score, K and λ are constants, lnK = natural logarithm of K and ln2 = natural logarithm of 2. In addition to substitution scores, raw score calculation also took into account affine gap penalties with a gap existence penalty = 11 and gap extension penalty = 1.

For this study, K = 0.041, λ = 0.267, m = length of query protein and total number of amino acids in fly proteome also known as n = entire length of fly proteome present in the reference database = 17,879,049,827, were used for E-value calculations. All the equations and parameters used here were taken from PSIBLAST results. For scoring 15-mer alignment, m = 15 and n = 8,629,350 were used. The value of n was calculated after taking into account all possible 15-mers centered on all lysines of fly proteome present in the reference database.

For every protein pair (namely human-fly and mouse-fly), there are 10 alignments from PSIBLAST (both the jobs yield 5 alignments each) results, including all the alignments that would have introduced bias in clustering analysis discussed later. Hence, for every protein pair the alignment with lowest re-calculated E-value was chosen for further analysis.

### 2.3 Frequent item-set mining based clustering of 15-mers centered on annotated SUMOylation sites

Frequent item-set mining methods are commonly used for finding patterns in customer transaction data in the retail industry, for example market basket analysis. Apriori algorithm is a commonly used method in the field of frequent item-set mining. In this study, Apriori algorithm [13] was used to detect commonly occurring amino acid patterns in the 15-mer data obtained from PSIBLAST analysis.

The human and mouse query proteins as well as the fly homologs identified in the respective PSIBLAST jobs have redundancy. This could introduce bias in the clustering analysis. Hence, all of these protein lists were culled using h-CD-HIT server (http://weizhong-lab.ucsd.edu/cdhit_suite/cgi-bin/index.cgi?cmd=h-cd-hit) [14–17] . The culling process was hierarchical in nature and was carried out with 3 different identity cutoffs – 90%, 60% and 30% respectively (the results discussed here were obtained at 30% redundancy). Details of non-redundant protein lists and 15-mers centered on SUMOylation sites present in these proteins are given below (Table-3).

**Table-3:** Summary of protein lists obtained from h-CD-HIT server

| **Query type** | **Protein type** | **Total list** | **nr (Non redundant list)** | **FASTA 15-mers (nr)** | **Aligned 15-mers (nr)** |
| --- | --- | --- | --- | --- | --- |
| Human | Human | 5283 | 3170 | 10245 | 10245 |
| Human | Fly | 8539 | 3725 | 8657 | 8657 |
| Mouse | Mouse | 468 | 373 | 549 | 549 |
| Mouse | Fly | 1700 | 707 | 876 | 876 |

It should be noted that for every fly homolog, only those 15-mers were chosen that occurred in longest alignments for the given fly protein. This was done because some fly proteins are aligned to multiple query proteins. Including 15-mers from all the alignments would have introduced repetitions that would have affected the clustering analysis described below.

At this point, it is important to define terminologies used in this analysis. An item is the frequency of each of the 20 amino acids to occur at every position in the 15-mer. Positions in a 15-mer are numbered from 0 to 14, the SUMOylated lysine is at 7th position. The 20 standard amino acids are sorted in an alphabetical order of their one letter code starting with Alanine (A) and ending with Tyrosine (Y). The SUMOylated lysine (7K) is omitted from the analysis since it is common to all 15- mers. Possible items could be 0A, 0C, 0D ….. 6V, 6W, 6Y and 8A, 8C, 8D ….. 14V, 14W, 14Y. In other words, there are 14 x 20 = 280 unique items in the data of a given 15-mer category. Another term used in this analysis is called support, which is frequency of a given item (or item-set) in the data of a given 15-mer category divided by total number of 15-mers of a given category. In other words, support is probability or normalized frequency. All 15-mers having gaps or occurring near N-terminus or C-terminus are excluded from this analysis.

The Apriori algorithm can be explained in the steps given below –

i. Calculate support values for each of the 280 items. Each item could be thought of as an item-set of size equal to 1. Select all size-1 item-sets that have their support greater than or equal to their support cutoff.
ii. Generate all possible new item-sets of size = 2.New item-sets are generated by extending item-sets from previous step by an item, such that the item is a member of an item-set from previous step.
iii. Support values are calculated for new item-sets. All item-sets with support greater than or equal to support cutoff are selected.
iv. This process of generation and selection of new item-sets having size greater by 1 than previous step, is continued till no new item-sets can be created. At this step, the algorithm ends.

The item-sets having size-2 resulting from the Apriori algorithm were processed further. In case a group of size-2 item-sets satisfy the following conditions, they are combined together and their support values are added up –

i. The given group of size-2 item-sets should have 1 common item and the varying item should have the same position but different amino acid.
ii. All the item-sets of the given group should have their support values greater than or equal to support cutoff for combining 2-mer item-set for the given 15-mer category in accordance to Table-4.

**Table-4:** Support cutoffs for each 15-mer category used as input to Apriori algorithm and support cutoff for clustering 2-item-sets for each 15-mer category

| **Query type** | **Protein type** | **Support cutoff Apriori (FASTA 15-mer) %** | **Support cutoff to cluster 2-mer (FASTA 15-mer) %** | **Support cutoff Apriori (Aligned 15-mer) %** | **Support cutoff to cluster 2-mer (Aligned 15-mer) %** |
| --- | --- | --- | --- | --- | --- |
| Human | Human | 0.425 | 1.5 | 0.425 | 1.3 |
| Human | Fly | 0.65 | 0.82 | 0.65 | 0.79 |
| Mouse | Mouse | 2.6 | 3 | 2.6 | 3 |
| Mouse | Fly | 0.85 | 1.5 | 0.85 | 1.5 |

Let us understand the combining exercise with an illustration. For example, consider 3 size-2 item-sets: 6I-9E, 6L-93 and 6V-9E. These item-sets have 1 common item namely 9E and the varying items – 6I, 6L and 6V have same position – 6 but varying amino acids namely – I, L and V. Since, the 2 conditions for combining size-2 item-sets has been met, the 3 item-sets will be combined into a consensus motif - 6[IVL]-9E, where I,V and L are the 3 amino acids that could occur at position 6 while E occurs at position 9. When the item-sets are combined into a consensus motif, their respective support values are added up and the sum is assigned to the consensus motif. Given below are support cutoffs used for different 15-mer categories (Table-4).

### 2.4 Hierarchical clustering of protein sequences

The results from h-CD-HIT server discussed in the previous section contain proteins clustered according to their sequence identities. Given below are details of clusters from each 15-mer category (Table-5).

**Table-5:**
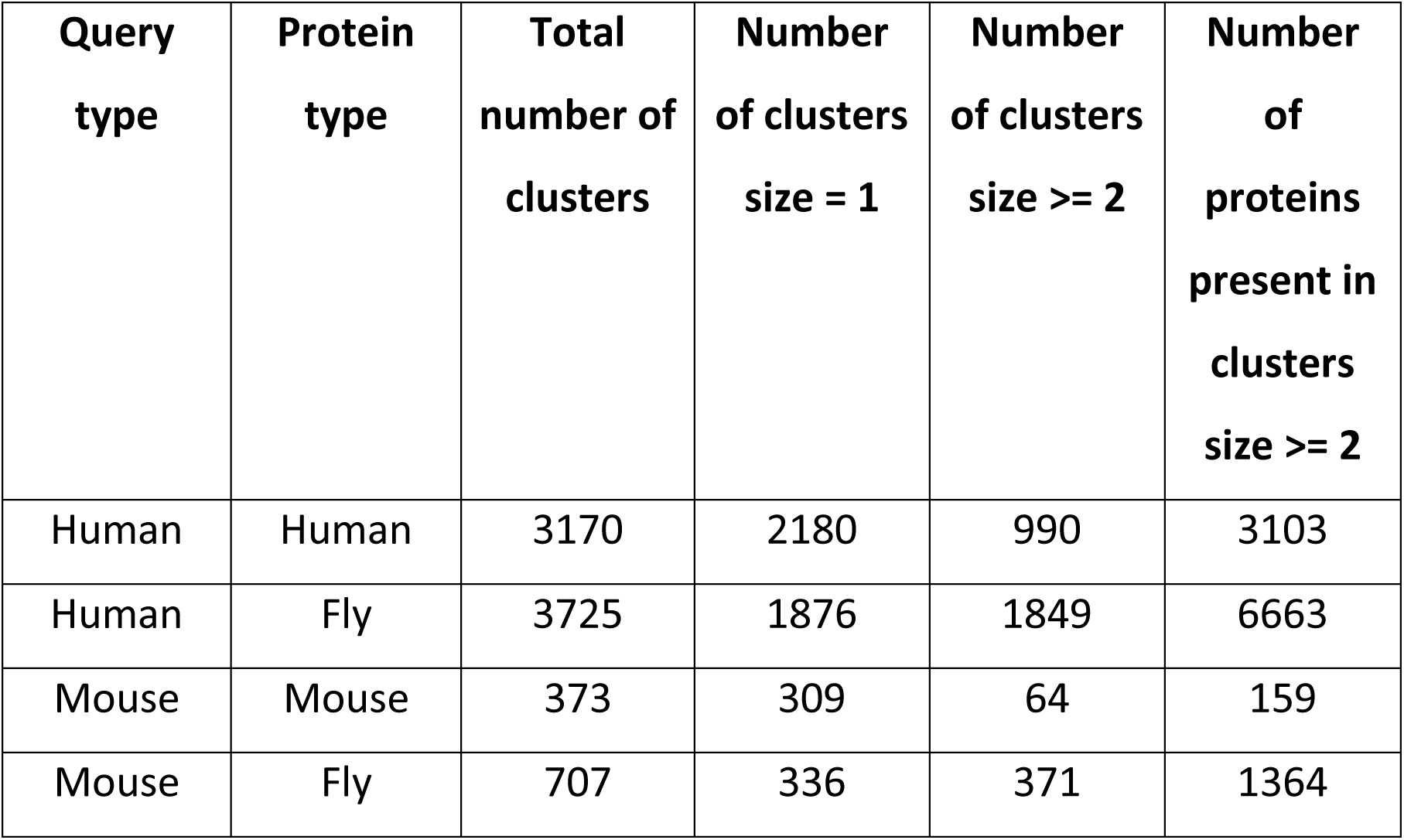
The Protein sequence clusters from h-CD-HIT server.

| **Query type** | **Protein type** | **Total number of clusters** | **Number of clusters size = 1** | **Number of clusters size $\geq 2$** | **Number of proteins present in clusters size $\geq 2$** |
| --- | --- | --- | --- | --- | --- |
| Human | Human | 3170 | 2180 | 990 | 3103 |
| Human | Fly | 3725 | 1876 | 1849 | 6663 |
| Mouse | Mouse | 373 | 309 | 64 | 159 |
| Mouse | Fly | 707 | 336 | 371 | 1364 |

Multiple sequence alignments were constructed for every cluster using SALIGN function in MODELLER 9.17 [18]. All SUMOylation sites that line up at the same position in the MSA were grouped together and 15-mers centered on these lysines were extracted (discussed later in Results section). These clusters of SUMOylation sites were sorted in descending order by their sizes. Motifs were extracted from these 15-mer clusters. In this case, a motif is a 15-mer sequence such that each of the 15 positions is represented by the most abundant amino acid at that position in the 15-mer cluster. In case a position has its most abundant amino acid having frequency less than 70%, then x is included at that position in the motif sequence. The frequency cutoff of 70% was used because any cutoff less than that would have been too lenient and it would have introduced noise in the motif sequence.

### 2.5 Clustering of Gene Ontology terms

This analysis was done to identify patterns related to protein functions for the identified homologs. Gene Ontology terms of every protein from both human and mouse PSIBLAST data were searched in the UniProt KB database. These terms can be divided into 3 categories - Cellular Component (GO C), Molecular Function (GO F) and Biological Process (GO P). It is possible for a protein to have one or more GO C, F and P terms associated with it. Each category of terms was clustered independently. All proteins that have the same term were grouped into the same cluster. For example, all proteins having Gene Ontology cellular component term as “nucleus” were grouped together into one cluster. Finally, all the clusters of a given category were sorted in descending order by their size.

### 2.6 Computational tools and programs used in this study

All the data extraction and analysis steps were carried out in Python version 2.7.5 and mathematical calculations were done using Numeric Python (NumPy) version

1.7.1 [19] respectively. Venn diagrams discussed later in the results section were plotted using VennDiagram library version 1.6.20 [20] in R version 3.4.4 [21] .

## 3 Results

### 3.1 Quantitative summary of results

#### 3.1.1 Predictions from human protein PSIBLAST

Given below is a summary of the results obtained from PSIBLAST-based analysis of 9256 human proteins described in [9] (Table-6). Out of a total 9256 proteins, about 5283 (57%) proteins have around 8539 fly orthologs. As for the 15-mers centered on SUMOylated lysines, 5283 human proteins contain 19823 15-mers centered on FASTA and aligned protein sequences. There around 52332 15-mers entered on annotated SUMOylated lysines from 8539 fly orthologs. There are about 4591 fly genes that encode for the 8539 fly proteins identified in this study. The CG-names and FlyBase FBgn identifiers for these genes were obtained from FlyBase release FB2018_05 (https://flybase.org/) [22] .

**Table-6:**
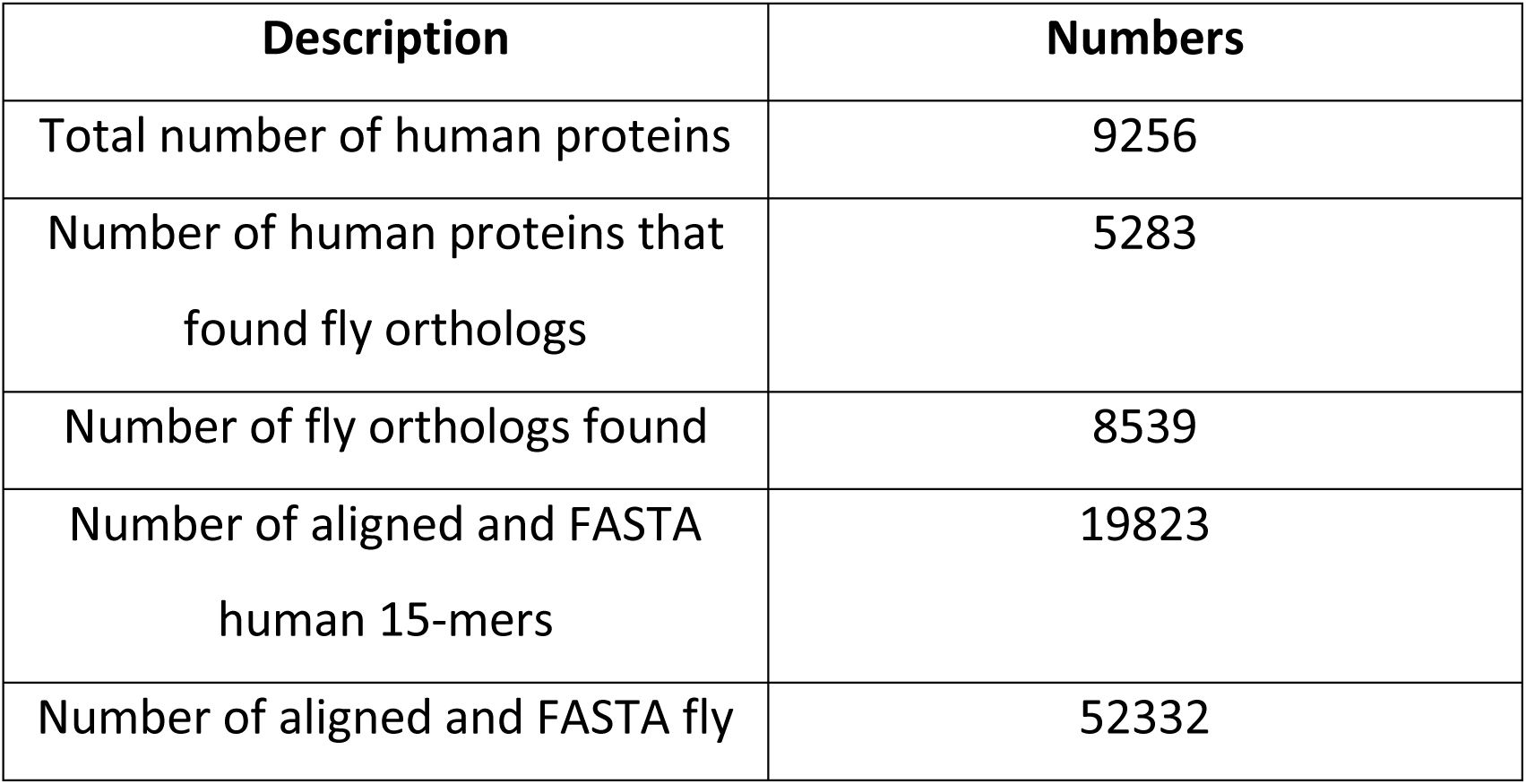
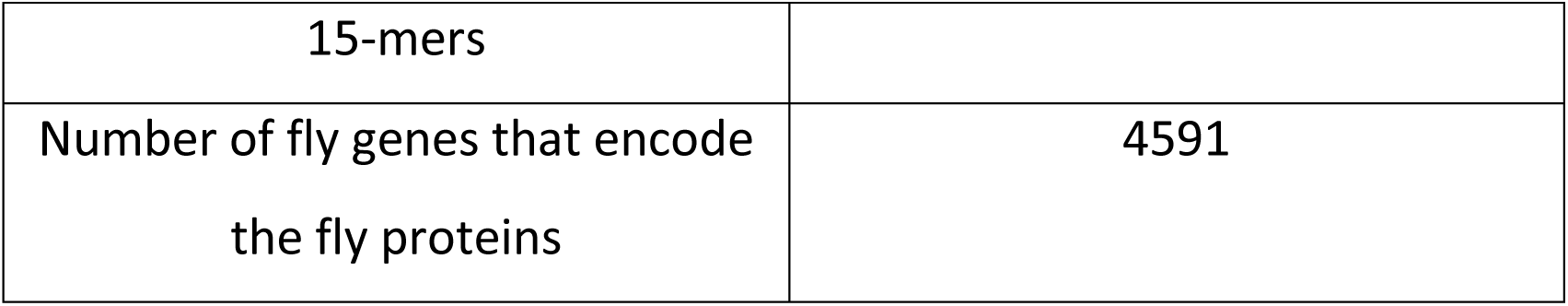
Summary of results obtained from PSIBLAST of human proteins

SUMOylated lysines are known to conform to a consensus sequence motif – ψ – K – x – (E/D) – where ψ = any aliphatic, hydrophobic amino acid such as I / V / L, K = SUMOylation site, x = any amino acid and E/D = either glutamate or aspartate residue. Since ψ and x are variable amino acids, only the K-x-E motif was checked in the FASTA 15-mers centered on SUMOylation sites. Given below is a summary of the proportion of human and fly SUMOylation sites that conform to the K– x– (E/D) motif (Table-7). The proportions of human and fly SUMOylation sites that do not conform to the consensus motif are 67% and 74% respectively.

**Table-7:** Summary of proportion of SUMOylation sites in human data that conform to K-x-(E/D) motif

| **Motif status** | **Proportion of human SUMOylation sites** | **Proportion of fly SUMOylation sites** |
| --- | --- | --- |
| K – x – (E/D) | 3128 (16 %) | 6292 (12 %) |
| (E/D) – x – K | 2820 (14 %) | 6225 (12 %) |
| (E/D) – x – K – x – (E/D) | 517 (3 %) | 1079 (2 %) |
| None | 13358 (67 %) | 38736 (74 %) |

A Venn diagram comparing the overlap between lists of CG-names for genes encoding fly SUMO targets identified by different SUMO proteomics studies can be seen below (Figure-2). These studies could identify fly SUMO targets but not the SUMOylated lysines due to experimental difficulties. The CG-name lists were derived from fly proteins identified by human PSIBLAST data, Nie et al 2009 [2], Handu et al 2015 [3], Pirone et al 2017 (S2R+ cell lines) and Pirone et al 2017 (transgenic flies) [4] respectively. Around 1069 (23 %) of CG-names for fly orthologs found from the human PSIBLAST data were confirmed by at least one of the other studies (Figure-2). These 1069 CG-names also account for 91% of the 1169 CG-names identified by 2 or more studies compared in the Venn diagram given below (Figure-2). There are 5 proteins common among all studies. These 5 proteins are actin-5C, Hsp68, RNP-107kd, 14-3-3 epsilon and 14-3-3 zeta.

**Figure 2:**
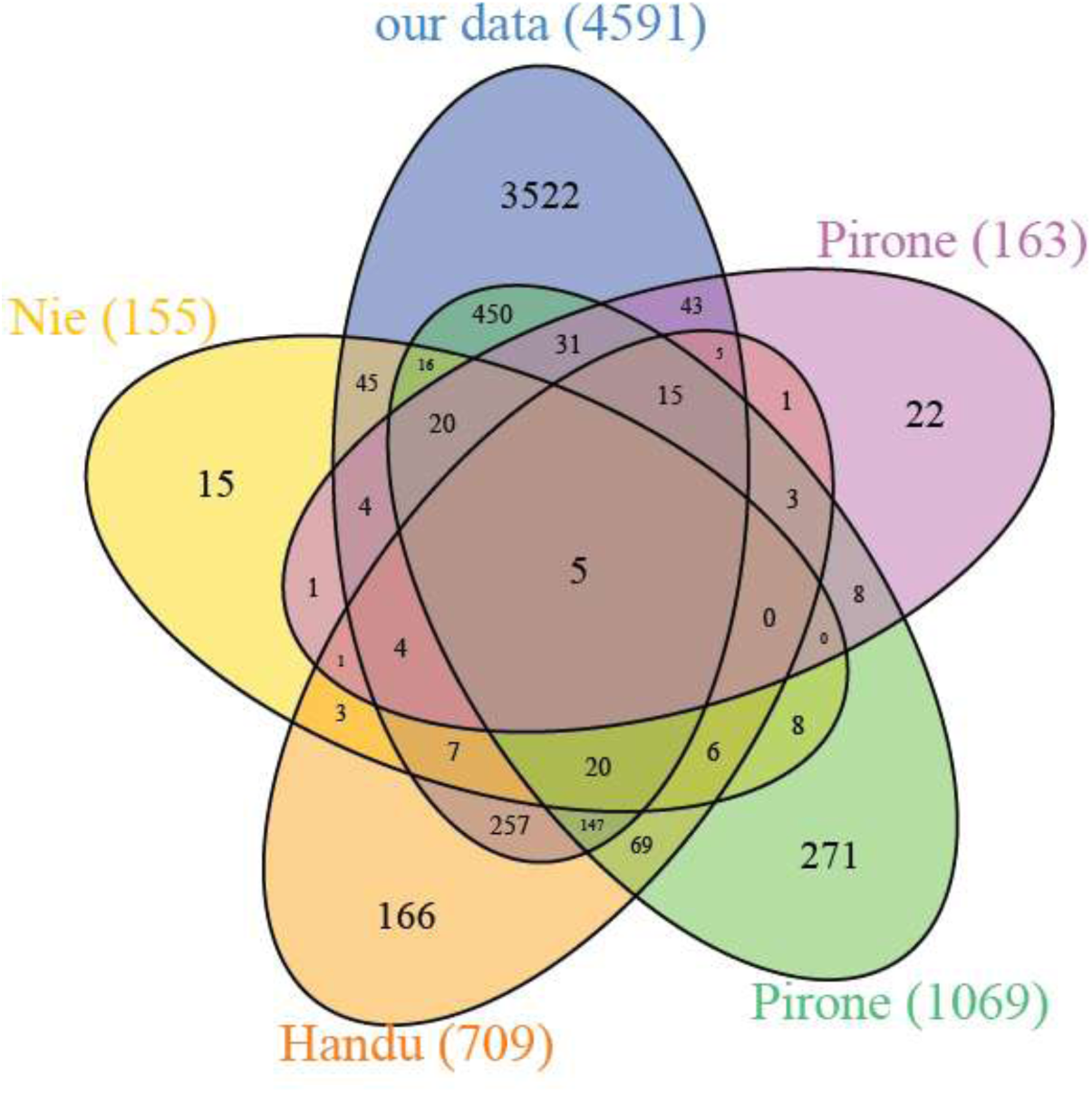
Venn diagram comparing gene CG-name list of fly orthologs from human PSIBLAST data with gene CG-name lists for fly SUMOylated proteins identified by other SUMO proteomics studies. Here, numbers in brackets indicate total number of CG-names for a given study and Nie – Nie et al 2009, Handu – Handu et al 2015, Pirone (1069) – Pirone et al 2017 data from S2R+ cell lines, Pirone (163) – Pirone et al 2017 data from transgenic flies.

Predictions from human PSIBLAST data also contain 3522 CG-names that were not identified by any of the 3 fly SUMO proteomics studies. 2194 of the 3522 fly genes encode at least one protein containing either K-x-(E/D), (E/D)-x-K or (E/D)-x-K-x-(E/D) consensus motifs. 1033 of the 3522 fly genes also encode proteins that have nucleus as their preferred cellular localization as indicated by their Gene Ontology Cellular Component terms. This observation is consistent with previous reports suggesting nucleus as a preferred cellular compartment of SUMOylated proteins [8, 9]. A detailed discussion of various Gene Ontology terms can be found later in this article.

#### 3.1.2 Predictions from mouse protein PSIBLAST

Given below is a summary of the results obtained from PSIBLAST of mouse proteins (Table-8). Out of 970 mouse proteins, around 468 proteins (48%) have 1700 fly orthologs. There are 769 SUMOylated lysines in 468 mouse proteins.

**Table-8:**
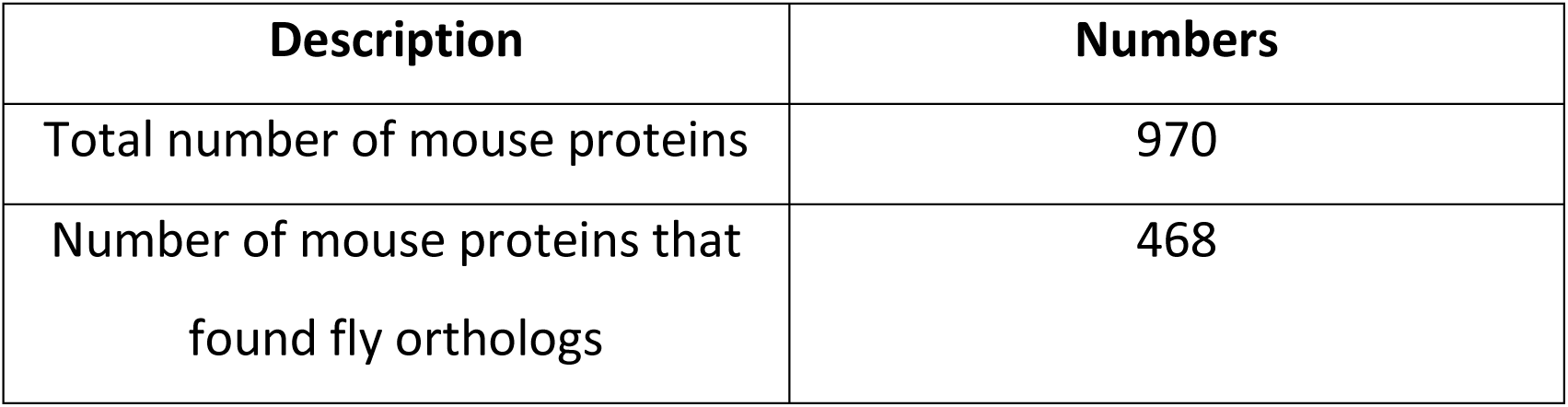
Summary of results obtained from PSIBLAST of mouse proteins

Similarly, the 1700 proteins contain 3700 annotated SUMOylated lysines. There are 936 fly genes that encode for the 1700 fly orthologs.

The proportions of K-x-(E/D) motif lysines in mouse and fly FASTA 15-mers are similar to the proportions reported for human PSIBLAST data in previous section (Table-9). Thus, around 57% of mouse SUMOylation sites and 76% of fly annotated SUMOylation sites do not conform to the K-x-(E/D) motif.

**Table-9:**
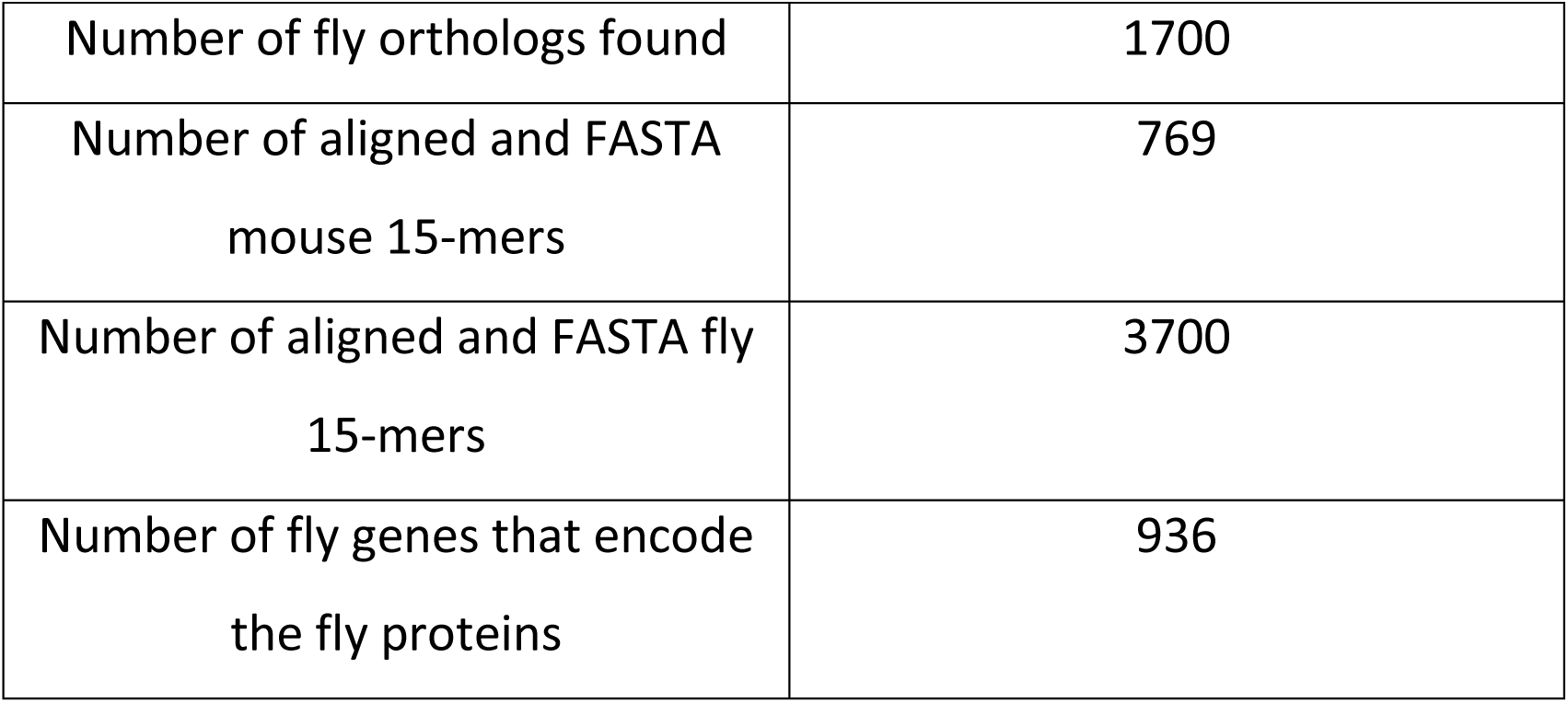
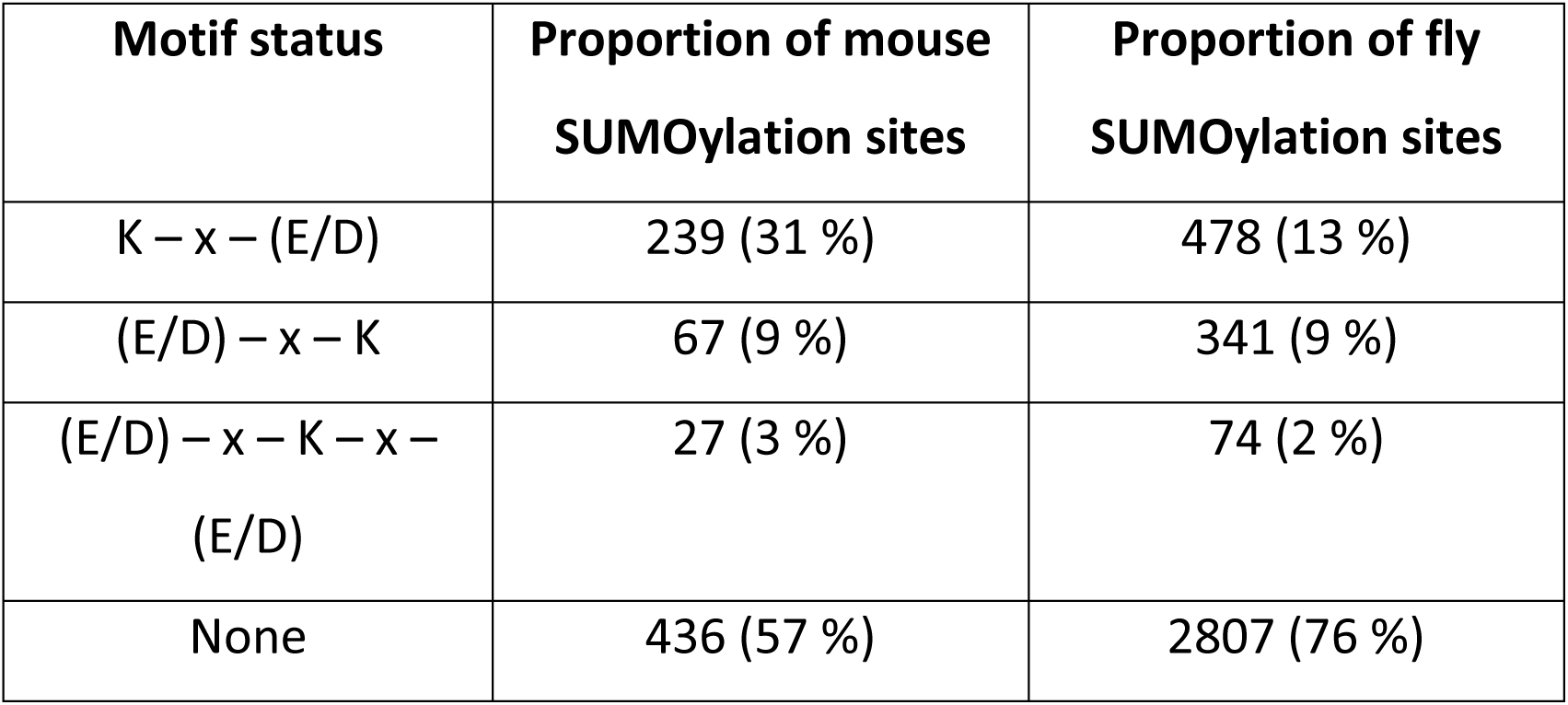
Summary of proportion of SUMOylation sites in mouse data that conform to K-x-(E/D) motif

Given above is a Venn diagram comparing overlap between CG-name lists of genes encoding fly orthologs identified from mouse PSIBLAST analysis and other SUMO proteomics studies (Figure-3A, Notations are the same as Figure-2).

**Figure 3:**
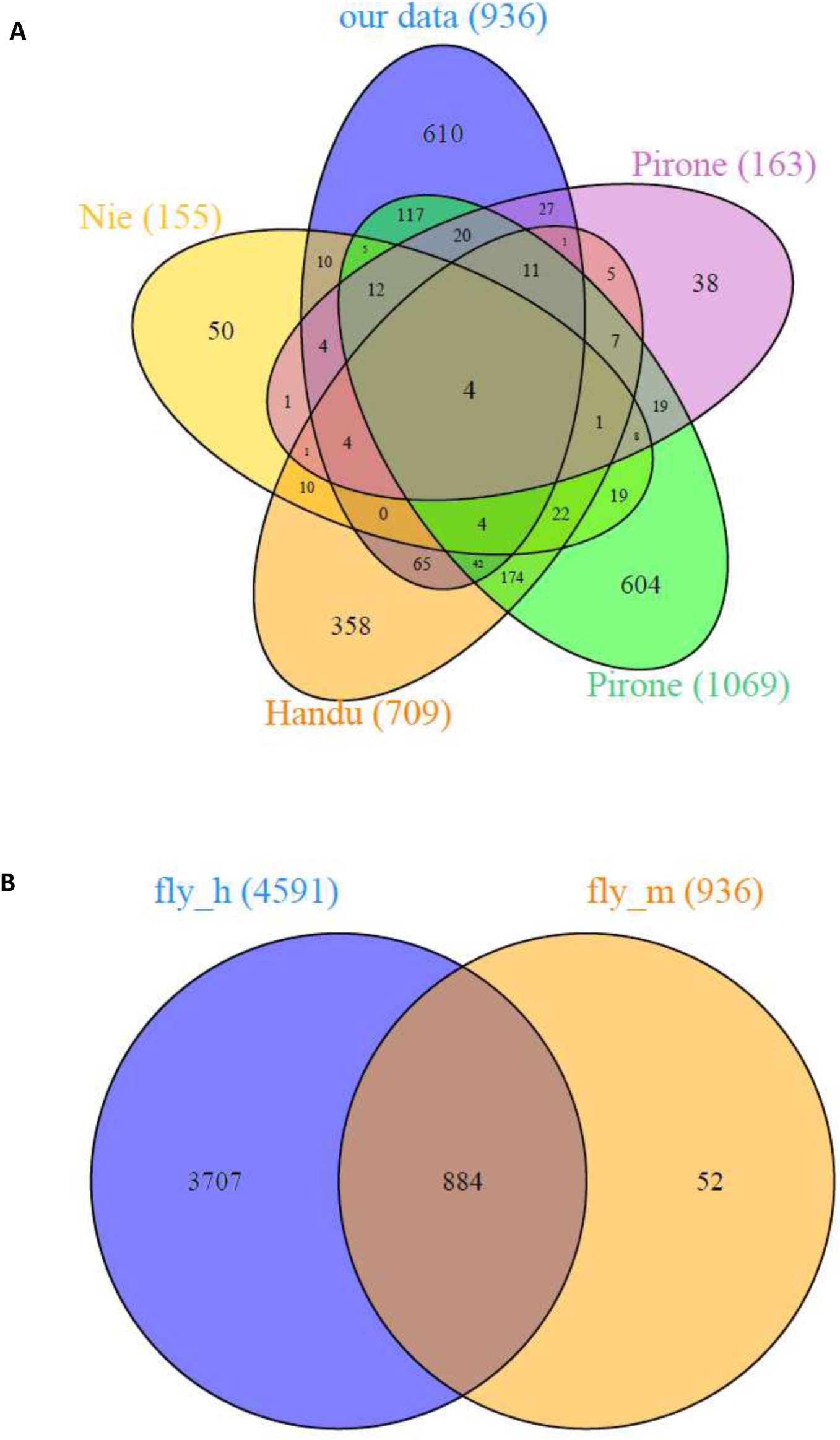
A: Venn diagram comparing gene CG-name list of fly orthologs identified from mouse PSIBLAST data with gene CG-name lists of fly SUMOylated proteins identified by other SUMO proteomics studies namely – Nie et al 2009, Handu et al 2015, Pirone et al 2017 (S2R+ cell lines) and Pirone et al 2017 (transgenic flies) Notations are same as Figure-2. B: Venn diagram comparing CG-name lists of fly orthologs identified by human PSIBLAST data and mouse PSIBLAST data.

Around 326 (35%) of CG-names for fly orthologs found from mouse PSIBLAST analysis were confirmed by at least one of the other studies. The 4 proteins common in all studies are RNP-107kd, actin-5C, 14-3-3 epsilon and 14-3-3 zeta. In addition, overlap between CG-name lists for genes encoding fly orthologs identified by human PSIBLAST data and mouse PSIBLAST data can be seen above (Figure-3B). Around 884 (94%) of CG-names from mouse data were confirmed by CG-names from human data.

### 3.2 Proteins identified by different fly SUMO proteomics experiments

The Venn diagram derived from human PSIBLAST data (Figure-2) consists of proteins that belong to different overlap categories. Overlap categories could range from 0 to 4, where either a protein was detected by none of the fly proteomics studies to as many as all 4 studies. Proteins detected by all 4 studies have been discussed in the previous section. For the sake of analysis, proteins could be divided into 4 different categories, 1 – detected by 3 of the 4 studies, 2 – detected by 2 of the 4 studies, 3 – detected by 1 of the 4 studies and 4 – detected uniquely in this study. All the 4 categories are a subset of human PSIBLAST data.

For each of these 4 categories, proteins were sorted in an ascending order according to the minimum E-value the given protein could achieve out of all the alignments involving the given protein. Given below are top 5 proteins from every category (Table-10).

**Table-10:** Top 5 proteins and their respective UniProt / UniRef90 IDs for every overlap category.

| **Rank** | **Proteins**<br>**found by 3**<br>**studies** | **Proteins**<br>**found by 2**<br>**studies** | **Proteins**<br>**found by 1**<br>**study** | **Proteins**<br>**unique to our**<br>**study** |
| --- | --- | --- | --- | --- |
| 1 | P0CG69 :<br>Polyubiquitin | P17210 :<br>Kinesin heavy<br>chain | A1ZAB5 :<br>Protein<br>clueless | X2JCN4 :<br>Kugelei,<br>isoform E |
| 2 | Q9TVM2 :<br>Exportin-1 | A4V1F9 :<br>Ubiquitin-63E,<br>isoform C | A0A0B4KEX0 :<br>Protein<br>clueless | P54358 : DNA<br>polymerase<br>delta catalytic<br>subunit |
| 3 | P13060 :<br>Elongation<br>factor 2 | Q59E34 : Mi-2, isoform B | Q9VPI9 :<br>LD20667p | Q7KU24 :<br>Chromodomai<br>n-helicase-DNA-binding<br>protein 1 |
| 4 | P15348 : DNA<br>topoisomeras<br>e 2 | E1JI46 : Mi-2,<br>isoform C | M9PIA6 : Mi-2, isoform D | UniRef90_Q8S<br>WV9 :<br>LD39323p<br>(Fragment) |
| 5 | P11147 :<br>Heat shock<br>70 kDa<br>protein<br>cognate 4 | Q9VF02 :<br>Helicase 89B,<br>isoform B | Q00174 :<br>Laminin<br>subunit alpha | P24014 :<br>Protein slit |

### 3.3 Motifs obtained from clustering analysis of 15-mer centered on SUMOylation sites

Shown below are top 5 motifs (in terms of support values) found in human PSIBLAST data using Apriori algorithm (Tables-11 and 12).

**Table-11:** Top 5 motifs found in FASTA and aligned 15-mers found in human proteins in human PSIBLAST data. Numbers in brackets indicate total number of motifs for the given category.

| Rank | Motif<br>human<br>FASTA 15-mer (5882) | Support<br>human<br>FASTA 15-mer (%) | Motif<br>human<br>aligned 15-mer (5885) | Support<br>human<br>aligned 15-mer (%) |
| --- | --- | --- | --- | --- |
| 1 | 6[ILV]-9E | 5.721 | 6[ILV]-9E | 5.621 |
| 2 | 9E-11[EL] | 3.538 | 9E-13[EKL] | 4.577 |
| 3 | 8E-9E | 2.013 | 2[EKL]-9E | 4.017 |
| 4 | 9E-12E | 1.904 | 9E-12[EK] | 3.26 |
| 5 | 4E-5E | 1.874 | 3[EL]-9E | 3.026 |

**Table-12:** Top 5 motifs found in FASTA and aligned 15-mers found in fly proteins in human PSIBLAST data. Numbers in brackets indicate total number of motifs for the given category.

| Rank | Motif fly<br>FASTA 15-mer (832) | Support fly<br>FASTA 15-mer (%) | Motif fly<br>aligned 15-mer (889) | Support fly<br>aligned 15-mer (%) |
| --- | --- | --- | --- | --- |
| 1 | 6[ILV]-9E | 2.994 | 4[AEKLRS]-6L | 5.272 |
| 2 | 9E-13[EKL] | 2.982 | 8L-13[AKLR] | 3.896 |
| 3 | 5[DEL]-6L | 2.829 | 6[AILV]-9E | 3.78 |
| 4 | 0[EKL]-8L | 2.805 | 9E-13[AEKL] | 3.78 |
| 5 | 4[AER]-6L | 2.793 | 0[EKLR]-8L | 3.737 |

As can be seen above, the most commonly occurring motif in human PIBLAST data is 6[IVL]-9E (Tables-11 and 12).

Given below are top 5 motifs (in terms of support values) found in mouse PSIBLAST data using Apriori algorithm (Tables-13 and 14).

**Table-13:** Top 5 motifs found in FASTA and aligned 15-mers found in mouse proteins in mouse PSIBLAST data. Numbers in brackets indicate total number of motifs for the given category.

| Rank | Motif<br>mouse | Support<br>mouse | Motif mouse<br>aligned 15- | Support<br>mouse |
| --- | --- | --- | --- | --- |

|  | **FASTA 15-**<br>**mer (39)** | **FASTA 15-**<br>**mer (%)** | **mer (28)** | **aligned 15-**<br>**mer (%)** |
| --- | --- | --- | --- | --- |
| 1 | 6[ILV]-9E | 21.642 | 6[ILV]-9E | 19.744 |
| 2 | 4[EKSV]-9E | 13.993 | 9E-12[DESTV] | 16.667 |
| 3 | 9E-13[EGP] | 13.06 | 4[AEKS]-9E | 14.103 |
| 4 | 9E-12[DES] | 12.5 | 9E-13[EGP] | 13.333 |
| 5 | 3[LPS]-9E | 11.754 | 3[LST]-9E | 11.538 |

**Table-14:**
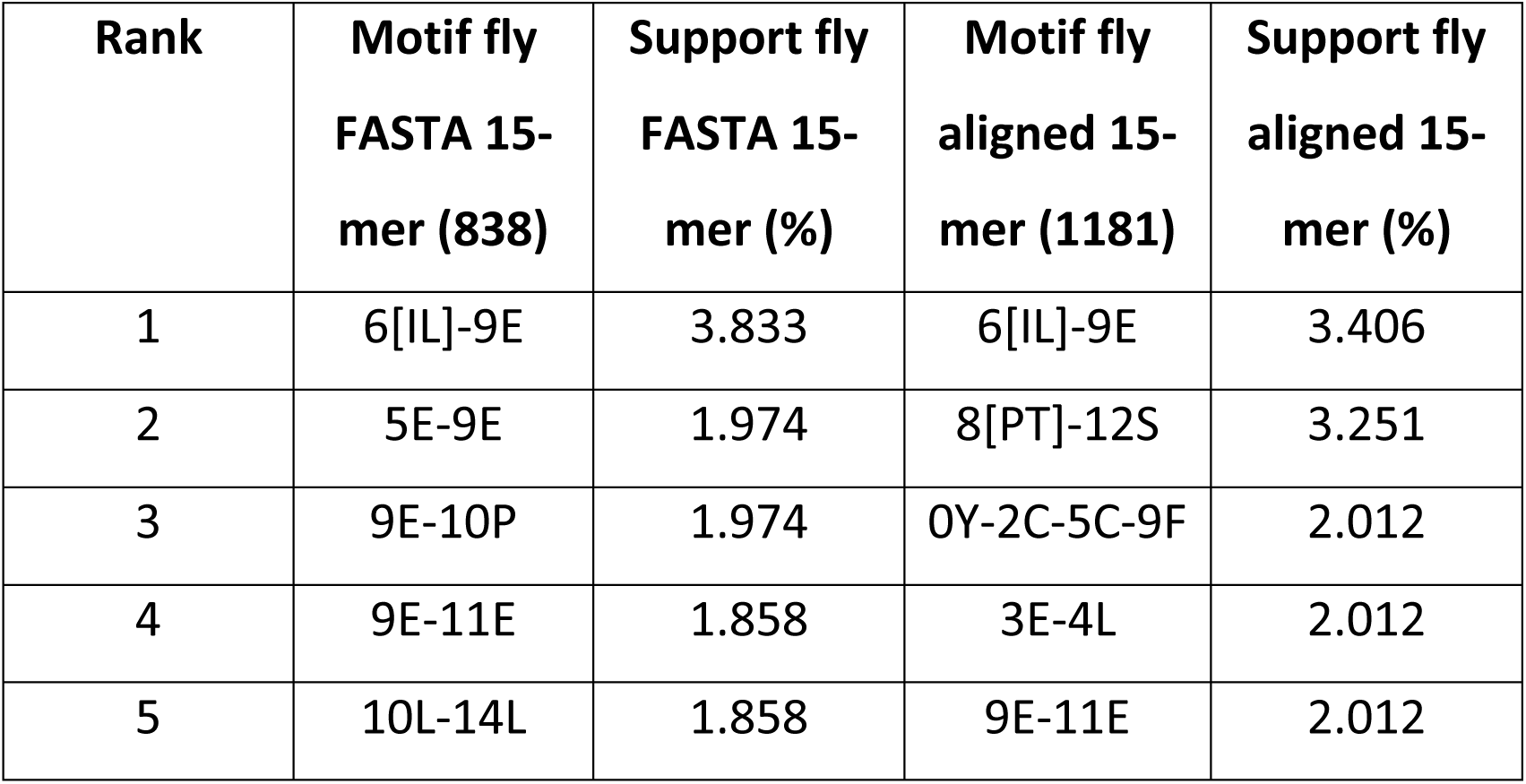
Top 5 motifs found in FASTA and aligned 15-mers found in fly proteins in mouse PSIBLAST data. Numbers in brackets indicate total number of motifs for the given category.

| **Rank** | **Motif fly**<br>**FASTA 15-**<br>**mer (838)** | **Support fly**<br>**FASTA 15-**<br>**mer (%)** | **Motif fly**<br>**aligned 15-**<br>**mer (1181)** | **Support fly**<br>**aligned 15-**<br>**mer (%)** |
| --- | --- | --- | --- | --- |
| 1 | 6[IL]-9E | 3.833 | 6[IL]-9E | 3.406 |
| 2 | 5E-9E | 1.974 | 8[PT]-12S | 3.251 |
| 3 | 9E-10P | 1.974 | 0Y-2C-5C-9F | 2.012 |
| 4 | 9E-11E | 1.858 | 3E-4L | 2.012 |
| 5 | 10L-14L | 1.858 | 9E-11E | 2.012 |

As can be seen above, 6[IVL]-9E is the amino acid motif with highest frequency in mouse PSIBLAST data (Tables-13 and 14).

All the motifs shown above (Tables-11 to 14) also contain the item 7K or central SUMOylated lysine. Recall from methods section that this item was omitted from input to Apriori algorithm because it occurs in all 15-mers. The item 7K should be considered while analyzing all the motifs (Tables-11 to 14). Thus, the motif 6[IVL]-9E can be expanded as 6[IVL]-7K-9E motif. In other words, this motif represents the SUMOylation consensus motif - ψ – K – x – (E/D) - where ψ is an aliphatic hydrophobic amino acid I, V or L occurring at 6^th^ position, 7K is the SUMOylated lysine and 9E indicates the glutamate residue occurring at the 9^th^ position in the 15-mer sequence.

### 3.4 Motifs obtained from clustering analysis of protein sequences

Given below are top 5 motifs (in terms of counts) occurring in MSAs built from proteins identified in human and mouse PSIBLAST data respectively (Tables-15 and 16).

While analyzing the tables given above, it is important to note that the count values are actually the total number of 15-mers in a given cluster. For an amino acid to be included in the motif sequence, it should have frequency greater than or equal to 70% of the count value at a given 15-mer position. Failing this criterion, a dummy amino acid x is added at that position in the motif sequence.

All the top 5 15-mer clusters from human proteins (Table-15) have the motif sequence HTGEKPYxCxxC. The MSA from zinc finger proteins contains 5 conserved repeats of the HTGEKPYxCxxC motif. Hence, there are 5 different instances of the same motif (Table-15). In addition to this motif, zinc finger proteins contain another motif sequence CxxCGKxF. However the CxxCGKxF motif occurs with a lower count as compared to HTGEKPYxCxxC motif.

**Table-15:**
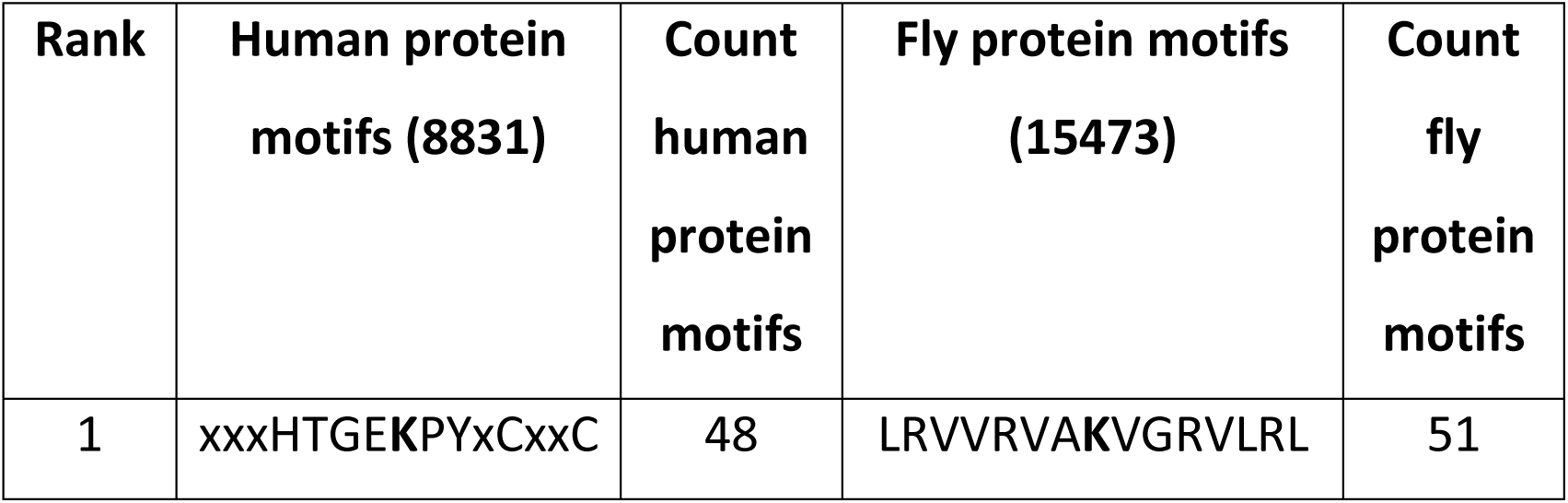
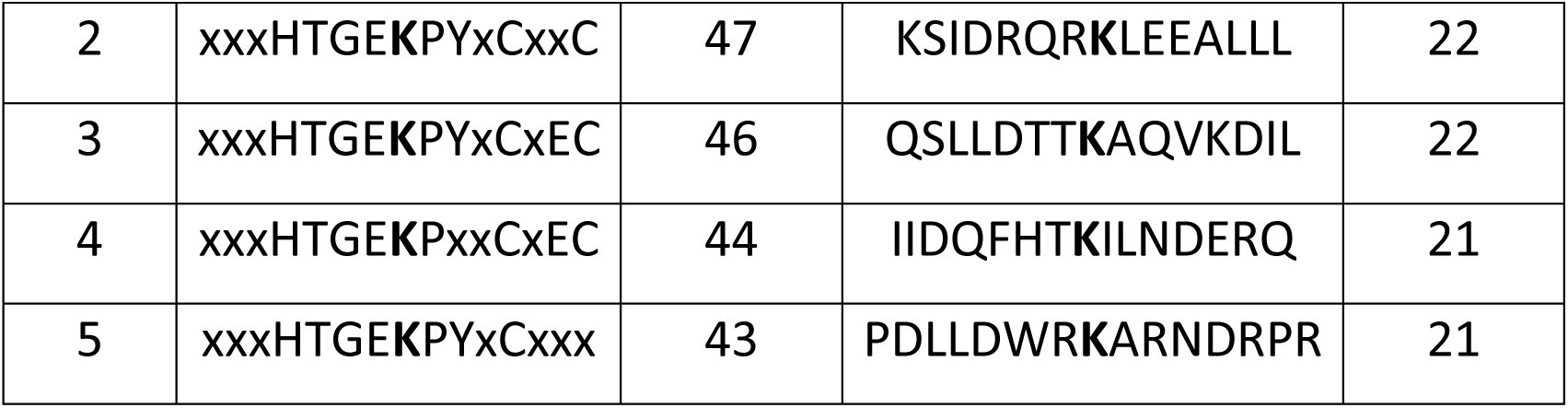
Top 5 motifs found in MSAs built from human and fly proteins in human PSIBLAST data. Numbers in brackets indicate total number of motifs for the given category.

Motif sequence with rank-1 from fly proteins (Table-15) occurs in MSA built from sodium channel proteins. Motifs ranked-2, 3, 4 and 5 occur in MSA built from short stop proteins. Similar to human proteins, fly proteins also contain zinc finger proteins and HTGEKPYxCxxC motifs albeit with a lower frequency.

Motifs ranked-1, 4 and 5 under mouse column (Table-16) occur in MSAs built from histones. Motif ranke-2 occurs in aldo-keto reductase family 1. Motif rank-3 occurs in chromobox proteins.

**Table-16:** Top 5 motifs found in MSAs built from mouse and fly proteins in mouse PSIBLAST data. Numbers in brackets indicate total number of motifs for the given category.

| Rank | Mouse protein motifs (266) | Count mouse protein motifs | Fly protein motifs (1041) | Count Fly protein motifs |
| --- | --- | --- | --- | --- |
| 1 | AAPAPxEKxPxKKKA | 4 | xEExxFPKATDxTFx | 17 |
| 2 | xxxxGLxKxxGxSNF | 3 | VxxxGxMKFxQxxxx | 17 |
| 3 | EADLVxAKEANxKCP | 3 | LEAFGNAKTVxNDNS | 17 |
| 4 | VSLxALKKxLAAXGY | 3 | xFxExSAKxxxNVxx | 16 |
| 5 | QAVLLPKKxxxxxxxx | 3 | xxLExQxKELxxxLx | 16 |

Motifs ranked-1, 2, 3 and 5 in fly columns (Table-16) occur in MSA from myosin heavy chain. Motif rank-4 occurs in Rab proteins.

### 3.5 New predictions made using motifs obtained from protein sequence clusters

A total of 13,922 out of 22,005 proteins of the fruit fly proteome were not picked up by either human or mouse PSIBLAST data. Hence, 15-mers centered on all lysine residues in these 13,922 proteins were extracted. Each of these 15-mers were scanned against a list of 11,227 motifs obtained after combining all fly motifs that had a count equal or greater than 2. The list of 11,227 motifs was created such that if one motif is a subset of another motif, then the larger motif was retained whereas the smaller motif was removed. This was done to ensure that the list of 11.227 motifs was non-repeating and unique in nature. For every query 15-mer the longest motif that matched the query 15-mer was found. This exercise led to identification of 2694 15-mers centered on new lysines from 1131 out of 13,922 proteins. These 2694 15-mers matched 1208 fly motifs. Given below are top 5 fly motifs that matched the most number of 15-mers centered on new lysines (Table-17).

**Table-17:** Top 5 fly motifs that matched most number of 15-mers centered on new lysines from remaining fly proteome. Numbers in brackets indicate total number of motifs for the given category.

| Rank | Fly motifs (1208) | New lysine count (2694) |
| --- | --- | --- |
| 1 | xVxxxxSKSxxxxxx | 151 |
| 2 | xxxxxxPKxxxNxKx | 60 |
| 3 | VxxxDxxKxxxVxxx | 55 |
| 4 | xxxxDxNKDxxxxxx | 48 |
| 5 | xHxxPxVKxxxxxxx | 42 |

Around 6 % of the lysines detected using motifs derived from protein MSAs conform to either forward or inverse orientation of the consensus motif (Table-18).

**Table-18:**
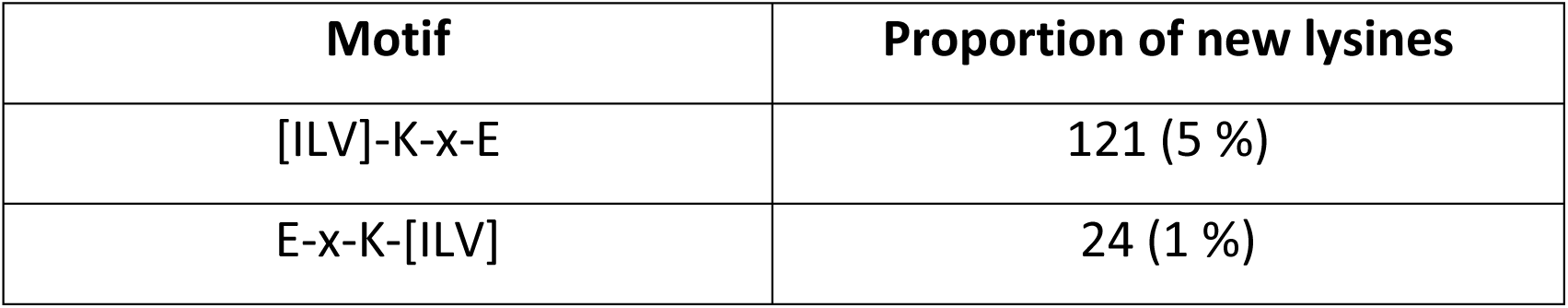
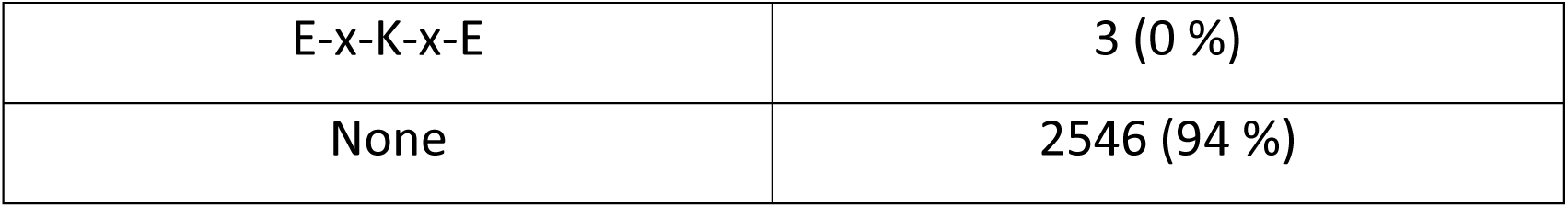
Summary of proportion of new lysines that conform to consensus motif.

The list of top 5 proteins shown in 2^nd^ column (Table-19) was obtained as follows. First, overlap (with respect to the 4 fly proteomics studies discussed earlier in Venn diagrams) was calculated for each of the 1131 proteins detected from motif matching exercise. Second, this list of 1131 proteins was sorted in descending order by their overlap values. From this sorted protein list, first 5 proteins that contain a lysine found using motif matching were chosen as long as the lysine conformed to either forward or inverse consensus motif.

**Table-19:** Top 5 proteins and their respective UniProt / UniRef90 IDs for every overlap category.

| Rank | Proteins found by motif matching (UniProt ID : protein name) | Overlap with other fly studies |
| --- | --- | --- |
| 1 | O97159 : Chromodomain-helicase-DNA-binding protein Mi-2 homolog | 2 |
| 2 | P36179 : Serine/threonine-protein phosphatase PP2A 65 kDa regulatory subunit | 2 |
| 3 | Q9NFP5 : SH3 domain-binding glutamic acid-rich protein homolog | 2 |
| 4 | Q9VPH7 : Eukaryotic peptide chain release factor subunit 1 | 2 |
| 5 | A0A0B4LH53 : N-ethylmaleimide-sensitive factor 2, isoform B | 1 |

### 3.6 Analysis of Gene Ontology terms

Given below is the summary of top 5 clusters obtained after analyzing Gene Ontology cellular component, molecular function and biological process terms from human PSIBLAST data (Tables-20 to 22).

**Table-20:** Top 5 GO C terms found in human and fly proteins in human PSIBLAST data. Numbers in brackets indicate total number of GO C terms for the given protein category.

| **Rank** | **GO C terms**<br>**human**<br>**proteins**<br>**(1206)** | **Count :**<br>**Proportion**<br>**%** | **GO C terms**<br>**fly proteins**<br>**(902)** | **Count :**<br>**Proportion**<br>**%** |
| --- | --- | --- | --- | --- |
| 1 | Nucleus | 2291 : 11% | Nucleus | 2333 : 14% |
| 2 | Nucleoplasm | 1886 : 9% | Cytoplasm | 1531 : 9% |
| 3 | Cytosol | 1724 : 8% | integral<br>component<br>of<br>membrane | 867 : 5% |
| 4 | Cytoplasm | 1324 : 6% | Cytosol | 703 : 4% |
| 5 | extracellular<br>exosome | 632 : 3% | plasma<br>membrane | 545 : 3% |

**Table-21:**
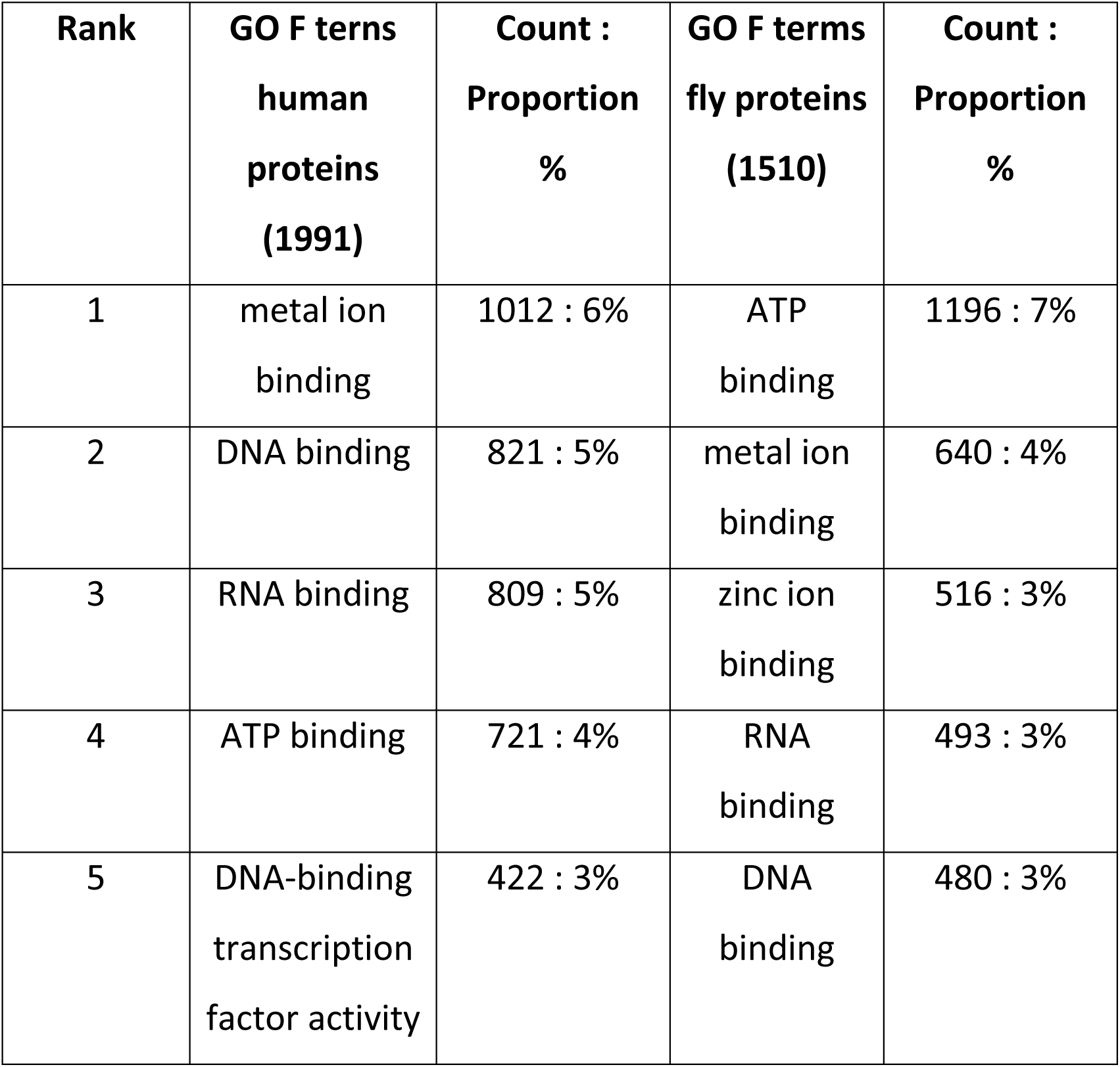
Top 5 GO F terms found in human and fly proteins in human PSIBLAST data. Numbers in brackets indicate total number of GO F terms for the given protein category.

| Rank | GO F terns<br>human<br>proteins<br>(1991) | Count :<br>Proportion<br>% | GO F terms<br>fly proteins<br>(1510) | Count :<br>Proportion<br>% |
| --- | --- | --- | --- | --- |
| 1 | metal ion<br>binding | 1012 : 6% | ATP<br>binding | 1196 : 7% |
| 2 | DNA binding | 821 : 5% | metal ion<br>binding | 640 : 4% |
| 3 | RNA binding | 809 : 5% | zinc ion<br>binding | 516 : 3% |
| 4 | ATP binding | 721 : 4% | RNA<br>binding | 493 : 3% |
| 5 | DNA-binding<br>transcription<br>factor activity | 422 : 3% | DNA<br>binding | 480 : 3% |

**Table-22:** Top 5 GO P terms found in human and fly proteins in human PSIBLAST data. Numbers in brackets indicate total number of GO P terms for the given protein category.

| Rank | GO P terms<br>human<br>proteins<br>(7037) | Count :<br>Proportion<br>% | GO P terms fly<br>proteins (3855) | Count :<br>Proportion<br>% |
| --- | --- | --- | --- | --- |
| 1 | regulation of<br>transcription,<br>DNA-templated | 455 : 1% | regulation of<br>transcription,<br>DNA-templated | 393 : 1% |
| 2 | positive<br>regulation of<br>transcription<br>by RNA<br>polymerase II | 441 : 1% | positive<br>regulation of<br>transcription by<br>RNA<br>polymerase II | 333 : 1% |
| 3 | negative<br>regulation of<br>transcription<br>by RNA<br>polymerase II | 392 : 1% | regulation of<br>transcription by<br>RNA<br>polymerase II | 289 : 1% |
| 4 | regulation of<br>transcription<br>by RNA<br>polymerase II | 283 : 1% | protein<br>phosphorylation | 240 : 1% |
| 5 | negative regulation of transcription, DNA-templated | 268 : 1% | negative regulation of transcription by RNA polymerase II | 237 : 1% |

As can be seen from the tables above (Tables-20 to 22), majority of the proteins from human PSIBLAST data localize to the nucleus. Most of these proteins bind DNA, RNA or ATP. Regulating transcriptional activity of RNA polymerase II seems to be the common biological process of these proteins.

Given below is the summary of top 5 clusters obtained after analyzing Gene Ontology cellular component, molecular function and biological process terms from mouse PSIBLAST data (Tables-23 to 25).

**Table-23:**
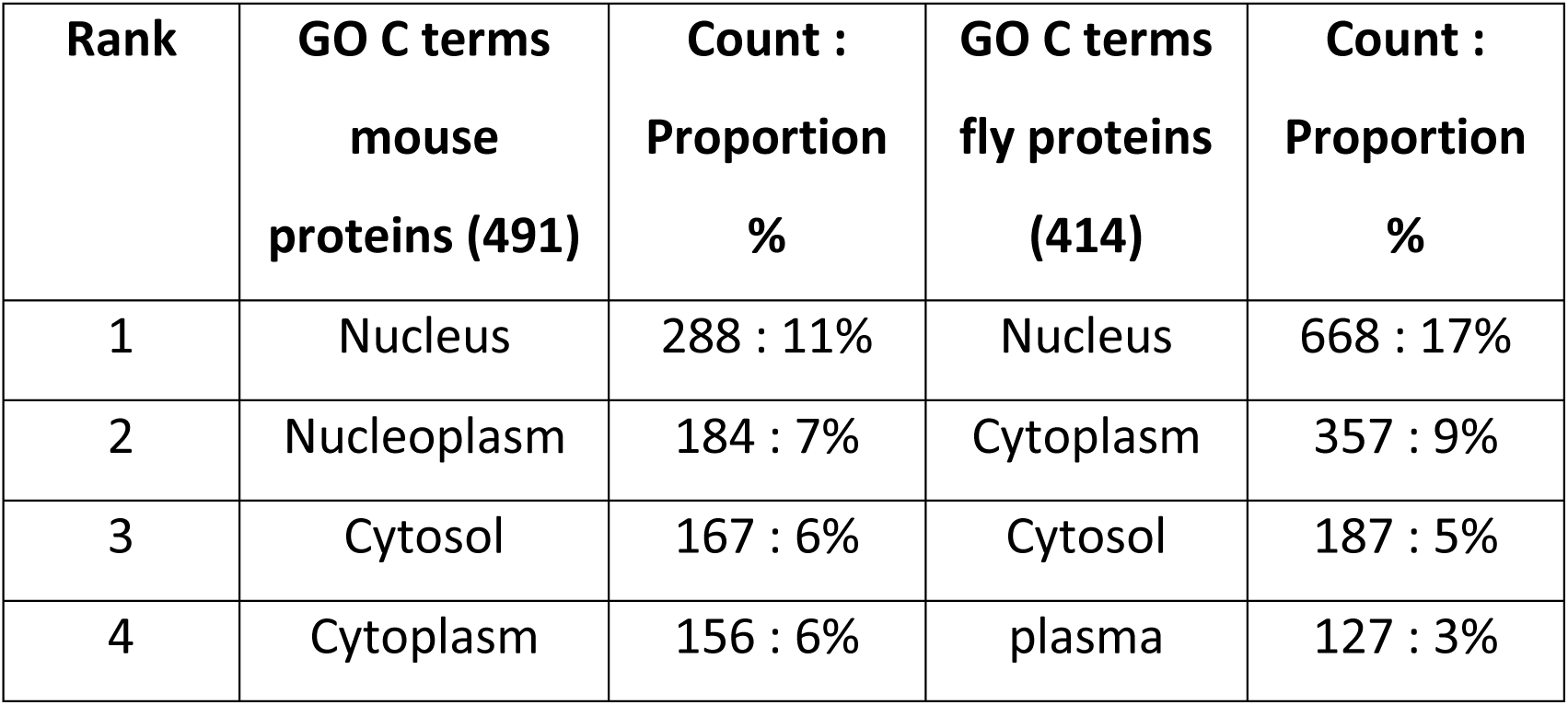
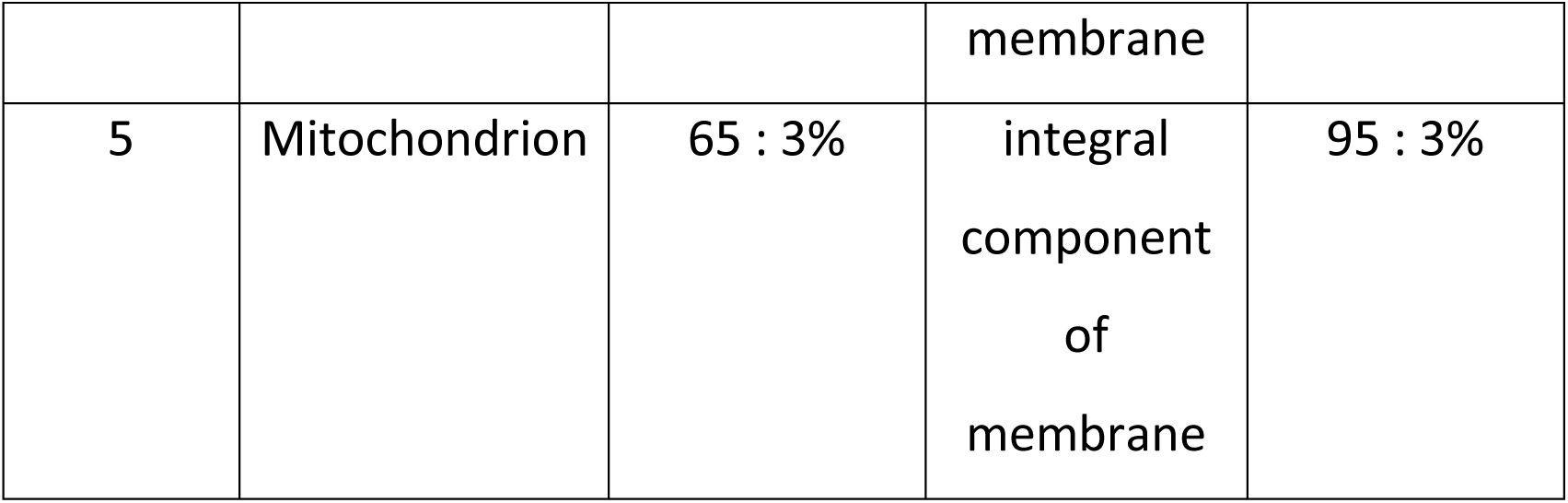
Top 5 GO C terms found in mouse and fly proteins in mouse PSIBLAST data. Numbers in brackets indicate total number of GO C terms for the given protein category.

**Table-24:**
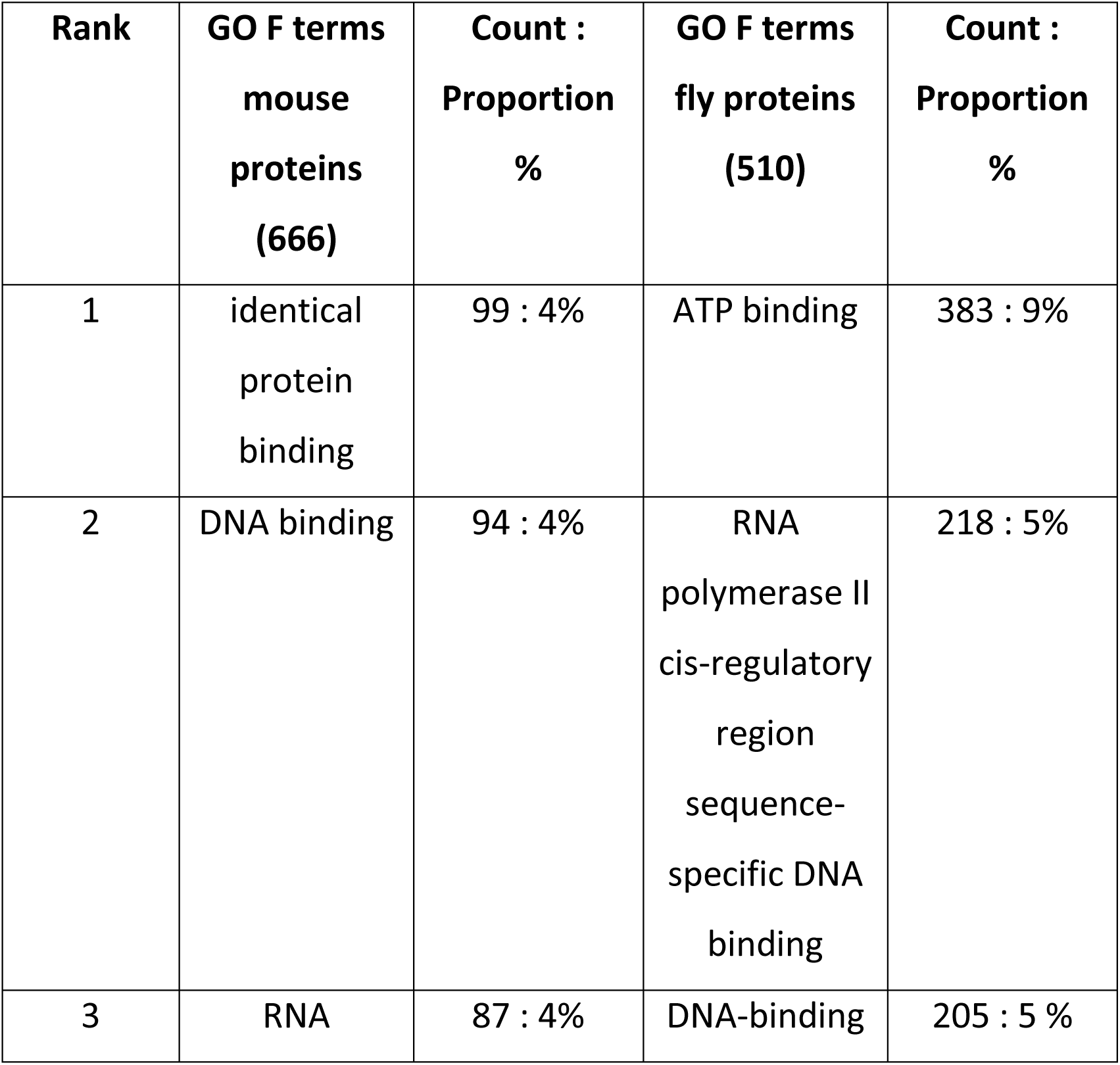
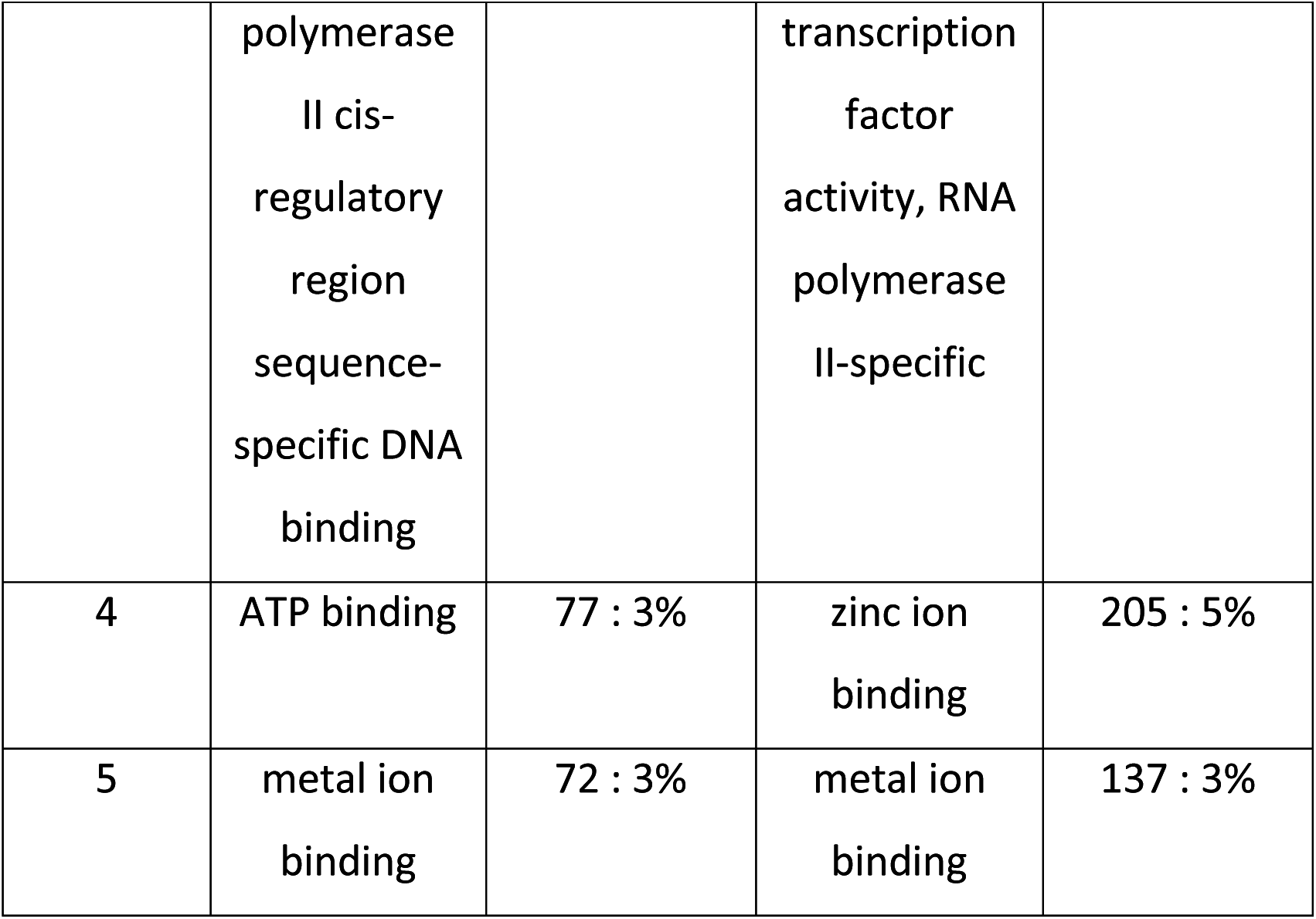
Top 5 GO F terms found in mouse and fly proteins in mouse PSIBLAST data. Numbers in brackets indicate total number of GO F terms for the given protein category.

**Table-25:**
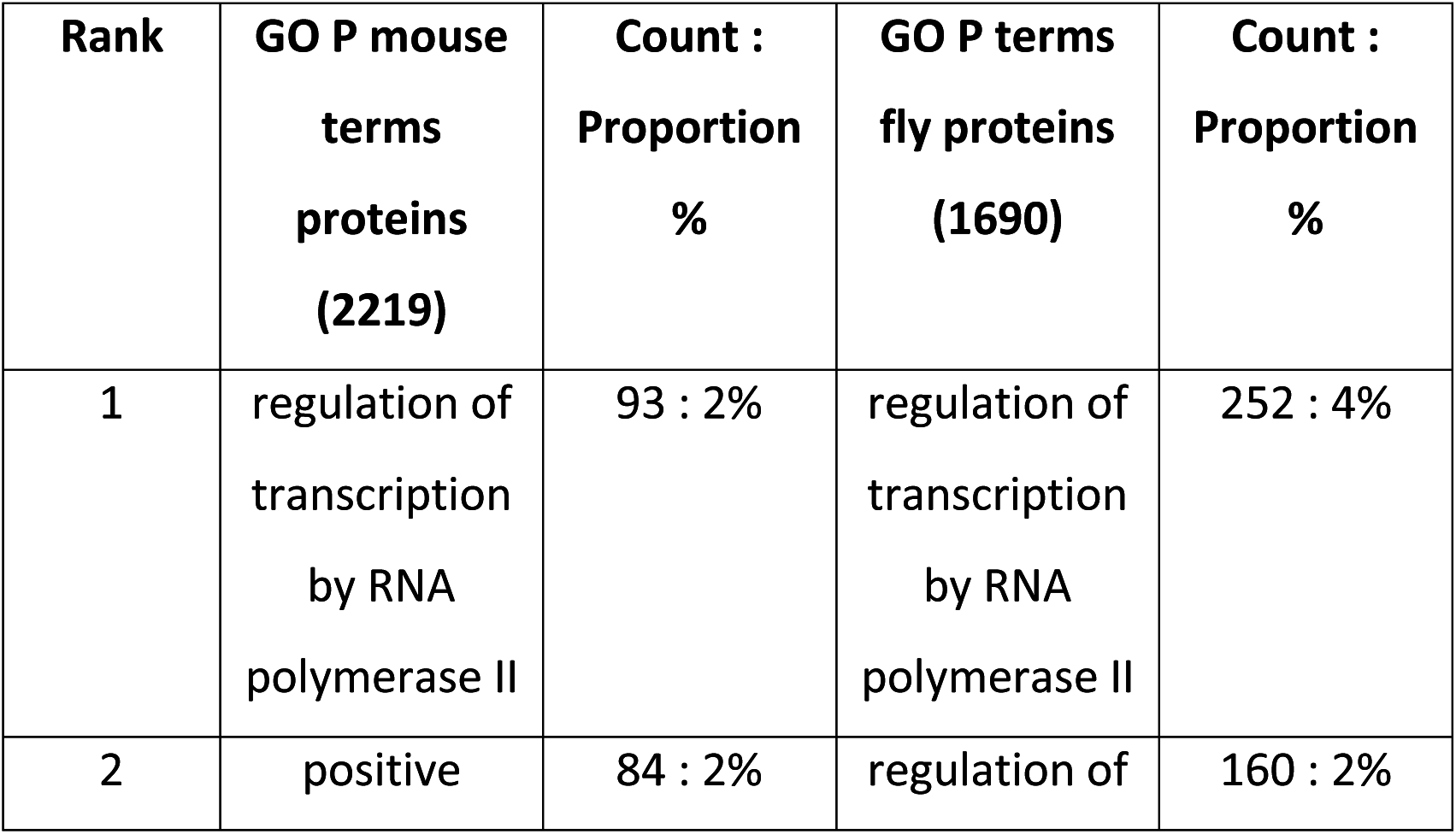
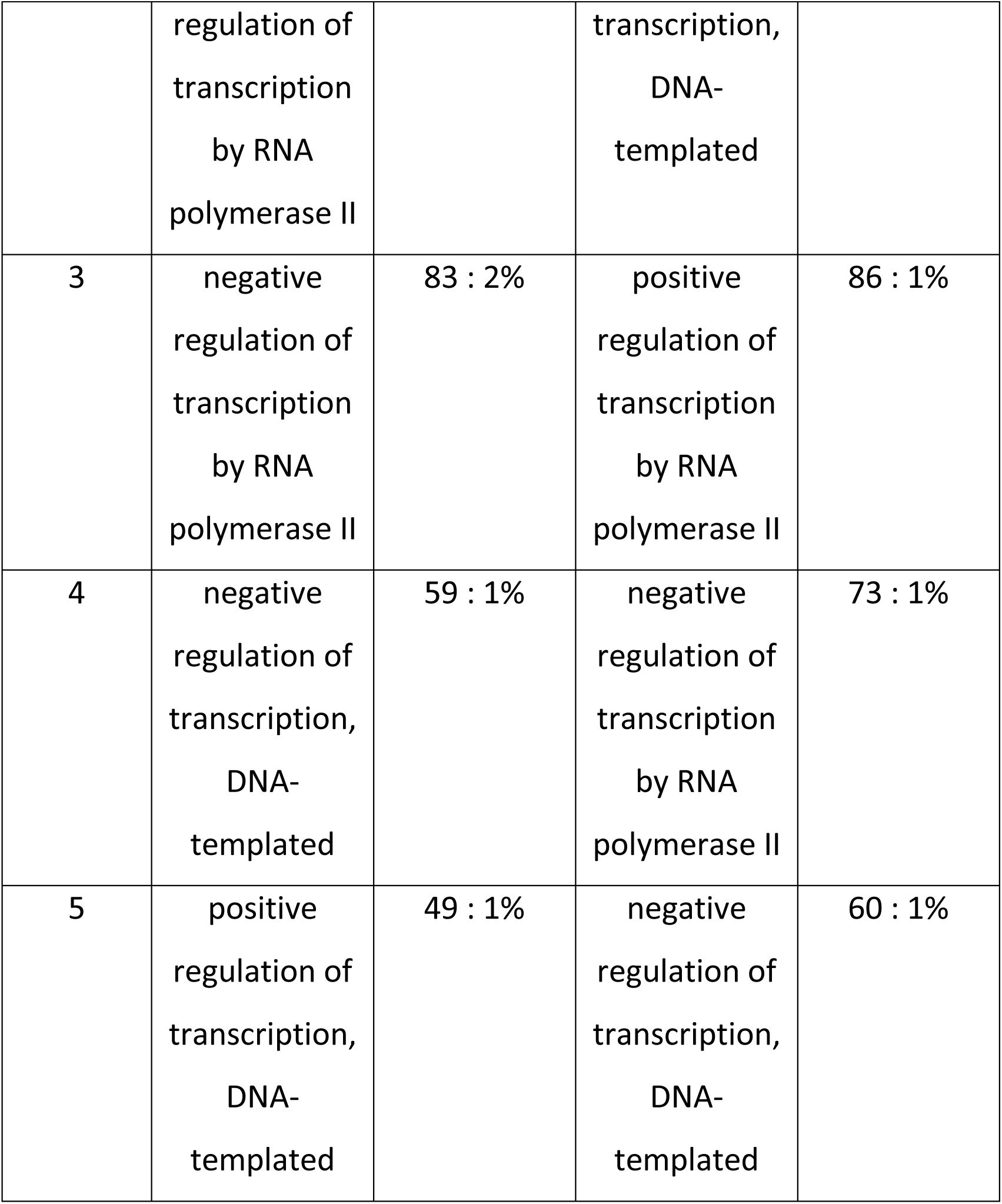
Top 5 GO P terms found in mouse and fly proteins in mouse PSIBLAST data. Numbers in brackets indicate total number of GO P terms for the given protein category.

As can be seen above (Tables-23 to 25), GO term analysis of mouse PSIBLAST data reveals that most of the proteins localize to the nucleus.

Most of these proteins bind metal ions, ATP, DNA or RNA. Regulating the transcriptional activity of RNA polymerase II seems to be the primary biological function of these proteins. Thus, taken together from GO term analysis of human and mouse PSIBLAST data, it seems that both categories of proteins have similar biological roles in a cellular environment.

### 3.7 Comparison between predictions from our study, GPS-SUMO and JASSA

The list of SUMOylated lysines predicted in this study was compared with the list obtained from sequence-based SUMOylation site prediction tools namely GPS-SUMO and JASSA. The fly proteins discussed in the present study could come from either fly proteome sequences or the UniRef90 database. Hence, all the 22,005 proteins of the fruit fly proteome and 1087 UniRef90 sequences were submitted as input to GPS-SUMO and JASSA. GPS-SUMO predicted 34,662 SUMOylation sites in 13,216 proteins from the fruit fly proteome and 3518 sites in 766 UniRef90 sequences. JASSA predicted 268,499 lysines in 21,034 proteins from the fruit fly proteome and 26,592 lysines in 1,077 UniRef90 sequences. Given below is a comparison between lysines predicted in this study and lysines predicted by GPS-SUMO and JASSA (Table-26).

**Table-26:** Overlap between lysines predicted in this study and predictions made by GPS-SUMO and JASSA.

|  | **Overlap of our study with results from GPS-SUMO** | **Overlap of our study with results from JASSA** | **Overlap between results from GPS-SUMO and JASSA** |
| --- | --- | --- | --- |
| Proteome | 2841 (8%) | 17,662 (7%) | 31,319 (90%) |
| UniRef90 | 363 (10%) | 2393 (9%) | 3181 (90%) |

The method presented in this study takes protein homology information into account. However, GPS-SUMO and JASSA do not consider homology information while predicting SUMOylation sites. This could be the reason for the low overlap between SUMOylated lysines predicted in this study and those predicted by GPS-SUMO and JASSA. Both GPS-SUMO and JASSA take local sequence environment around lysine residues into account while making predictions and prefer lysines conforming to consensus motifs. Hence, predictions made by both the tools are 90% similar.

## 4 Discussion

The work done in this study uses the concept of sequence homology for annotating SUMOylation sites in the proteome of fruit fly *Drosophila melanogaster* using information derived from human and mouse cells. Future studies could extend this idea to other post-translational modifications such as phosphorylation, ubiquitination, acetylation and others.

Clustering methods discussed in this study provide three kinds of information. First - amino acid preferences from a local sequence environment provided by 15- mer clustering exercise. This analysis confirmed the importance of the ψ–K-x-(E/D) consensus motif. Second - protein family specific preferences provided by protein sequence clustering exercise. This analysis showed that members of many protein families get SUMOylated, notable examples include zinc finger proteins, sodium channel proteins, histones and chromobox containing proteins. Third – cellular and molecular function preferences provided by GO term clustering exercise. This analysis showed that most of the SUMOylated proteins localize to nucleus, bind DNA / RNA and are involved in regulation of transcriptional activity.

The input information for this study was obtained from mass spectrometry-coupled proteomics experiments. The result of this *in silico* study can be experimentally validated in the near future. Such a combination of computational and experimental methods could be used for a better understanding of post-translational modifications in the near future.

## 6. Supplementary Data

All the raw data relevant to the present study can be found on Github here - https://github.com/yogendra-bioinfo/homology-based-SUMOylation-prediction. The significance of each data file can be found in a file named README.txt.

## References

[1] A. Flotho, F. Melchior, Sumoylation : A Regulatory Protein Modification in Health and Disease, (n.d.). https://doi.org/10.1146/annurev-biochem-061909-093311.

[2] M. Nie, Y. Xie, J.A. Loo, A.J. Courey, Genetic and proteomic evidence for roles of Drosophila SUMO in cell cycle control, Ras signaling, and early pattern formation, PLoS One. 4 (2009). https://doi.org/10.1371/journal.pone.0005905.

[3] M. Handu, B. Kaduskar, R. Ravindranathan, A. Soory, R. Giri, V.B. Elango, H. Gowda, G.S. Ratnaparkhi, SUMO-enriched proteome for drosophila innate immune response, G3 Genes, Genomes, Genet. 5 (2015) 2137–2154. https://doi.org/10.1534/g3.115.020958.

[4] L. Pirone, W. Xolalpa, J.O. Sigursson, J. Ramirez, C. Pérez, M. González, A.R. De Sabando, F. Elortza, M.S. Rodriguez, U. Mayor, J. V. Olsen, R. Barrio, J.D. Sutherland, A comprehensive platform for the analysis of ubiquitin-like protein modifications using in vivo biotinylation, Sci. Rep. 7 (2017) 1–17. https://doi.org/10.1038/srep40756.

[5] Q. Zhao, Y. Xie, Y. Zheng, S. Jiang, W. Liu, W. Mu, Z. Liu, Y. Zhao, Y. Xue, J. Ren, GPS-SUMO: A tool for the prediction of sumoylation sites and SUMO-interaction motifs, Nucleic Acids Res. 42 (2014) 325–330. https://doi.org/10.1093/nar/gku383.

[6] J. Ren, X. Gao, C. Jin, M. Zhu, X. Wang, A. Shaw, L. Wen, X. Yao, Y. Xue, Systematic study of protein sumoylation: Development of a site-specific predictor of SUMOsp 2.0, Proteomics. 9 (2009) 3409–3412. https://doi.org/10.1002/pmic.200800646.

[7] G. Beauclair, A. Bridier-Nahmias, J.-F. Zagury, A. Saïb, A. Zamborlini, JASSA: a comprehensive tool for prediction of SUMOylation sites and SIMs., Bioinformatics. 31 (2015) 3483–3491. https://doi.org/10.1093/bioinformatics/btv403.

[8] I.A. Hendriks, A.C.O. Vertegaal, A comprehensive compilation of SUMO proteomics, Nat. Rev. Mol. Cell Biol. 17 (2016) 581–595. https://doi.org/10.1038/nrm.2016.81.

[9] I.A. Hendriks, D. Lyon, D. Su, N.H. Skotte, J.A. Daniel, L.J. Jensen, M.L. Nielsen, Site-specific characterization of endogenous SUMOylation across species and organs, Nat. Commun. 9 (2018). https://doi.org/10.1038/s41467-018-04957-4.

[10] A. Bateman, M.J. Martin, S. Orchard, M. Magrane, R. Agivetova, S. Ahmad, E. Alpi, E.H. Bowler-Barnett, R. Britto, B. Bursteinas, H. Bye-A-Jee, R. Coetzee, A. Cukura, A. Da Silva, P. Denny, T. Dogan, T.G. Ebenezer, J. Fan, L.G. Castro, P. Garmiri, G. Georghiou, L. Gonzales, E. Hatton-Ellis, A. Hussein, A. Ignatchenko, G. Insana, R. Ishtiaq, P. Jokinen, V. Joshi, D. Jyothi, A. Lock, R. Lopez, A. Luciani, J. Luo, Y. Lussi, A. MacDougall, F. Madeira, M. Mahmoudy, M. Menchi, A. Mishra, K. Moulang, A. Nightingale, C.S. Oliveira, S. Pundir, G. Qi, S. Raj, D. Rice, M.R. Lopez, R. Saidi, J. Sampson, T. Sawford, E. Speretta, E. Turner, N. Tyagi, P. Vasudev, V. Volynkin, K. Warner, X. Watkins, R. Zaru, H. Zellner, A. Bridge, S. Poux, N. Redaschi, L. Aimo, G. Argoud-Puy, A. Auchincloss, K. Axelsen, P. Bansal, D. Baratin, M.C. Blatter, J. Bolleman, E. Boutet, L. Breuza, C. Casals-Casas, E. de Castro, K.C. Echioukh, E. Coudert, B. Cuche, M. Doche, D. Dornevil, A. Estreicher, M.L. Famiglietti, M. Feuermann, E. Gasteiger, S. Gehant, V. Gerritsen, A. Gos, N. Gruaz-Gumowski, U. Hinz, C. Hulo, N. Hyka-Nouspikel, F. Jungo, G. Keller, A. Kerhornou, V. Lara, P. Le Mercier, D. Lieberherr, T. Lombardot, X. Martin, P. Masson, A. Morgat, T.B. Neto, S. Paesano, I. Pedruzzi, S. Pilbout, L. Pourcel, M. Pozzato, M. Pruess, C. Rivoire, C. Sigrist, K. Sonesson, A. Stutz, S. Sundaram, M. Tognolli, L. Verbregue, C.H. Wu, C.N. Arighi, L. Arminski, C. Chen, Y. Chen, J.S. Garavelli, H. Huang, K. Laiho, P. McGarvey, D.A. Natale, K. Ross, C.R. Vinayaka, Q. Wang, Y. Wang, L.S. Yeh, J. Zhang, UniProt: The universal protein knowledgebase in 2021, Nucleic Acids Res. 49 (2021) D480–D489. https://doi.org/10.1093/nar/gkaa1100.

[11] S.F. Altschul, T.L. Madden, A.A. Schäffer, J. Zhang, Z. Zhang, W. Miller, D.J. Lipman, Gapped BLAST and PSI-BLAST: a new generation of protein database search programs., Nucleic Acids Res. 25 (1997) 3389–3402. https://doi.org/10.1093/nar/25.17.3389.

[12] A.A. Schäffer, L. Aravind, T.L. Madden, S. Shavirin, J.L. Spouge, Y.I. Wolf, E. V. Koonin, S.F. Altschul, Improving the accuracy of PSI-BLAST protein database searches with composition-based statistics and other refinements, Nucleic Acids Res. 29 (2001) 2994–3005. https://doi.org/10.1093/nar/29.14.2994.

[13] R. Agrawal, R. Srikant, Fast Algorithms for Mining Association Rules in Large Databases, in: Proc. 20th Int. Conf. Very Large Data Bases, Morgan Kaufmann Publishers Inc., San Francisco, CA, USA, 1994: pp. 487–499. http://dl.acm.org/citation.cfm?id=645920.672836.

[14] Y. Huang, B. Niu, Y. Gao, L. Fu, W. Li, CD-HIT Suite: a web server for clustering and comparing biological sequences, Bioinformatics. 26 (2010) 680–682. https://doi.org/10.1093/bioinformatics/btq003.

[15] W. Li, L. Jaroszewski, A. Godzik, Tolerating some redundancy significantly speeds up clustering of large protein databases, Bioinformatics. 18 (2002) 77–82. https://doi.org/10.1093/bioinformatics/18.1.77.

[16] W. Li, A. Godzik, Cd-hit: a fast program for clustering and comparing large sets of protein or nucleotide sequences, Bioinformatics. 22 (2006) 1658– 1659. https://doi.org/10.1093/bioinformatics/btl158.

[17] W. Li, L. Jaroszewski, A. Godzik, Clustering of highly homologous sequences to reduce the size of large protein databases, Bioinformatics. 17 (2001) 282–283. https://doi.org/10.1093/bioinformatics/17.3.282.

[18] A. Sali, T.L. Blundell, Comparative protein modelling by satisfaction of spatial restraints, J. Mol. Biol. 234 (1993) 779–815. https://doi.org/10.1006/jmbi.1993.1626.

[19] C.R. Harris, K.J. Millman, S.J. van der Walt, R. Gommers, P. Virtanen, D. Cournapeau, E. Wieser, J. Taylor, S. Berg, N.J. Smith, R. Kern, M. Picus, S. Hoyer, M.H. van Kerkwijk, M. Brett, A. Haldane, J.F. del Río, M. Wiebe, P. Peterson, P. Gérard-Marchant, K. Sheppard, T. Reddy, W. Weckesser, H. Abbasi, C. Gohlke, T.E. Oliphant, Array programming with {NumPy}, Nature. 585 (2020) 357–362. https://doi.org/10.1038/s41586-020-2649-2.

[20] H. Chen, P.C. Boutros, VennDiagram: a package for the generation of highly-customizable Venn and Euler diagrams in R, BMC Bioinformatics. 12 (2011) 35. https://doi.org/10.1186/1471-2105-12-35.

[21] R Core Team, R: A Language and Environment for Statistical Computing, (2018). https://www.r-project.org/.

[22] A. Larkin, S.J. Marygold, G. Antonazzo, H. Attrill, G. dos Santos, P. V Garapati, J.L. Goodman, L.S. Gramates, G. Millburn, V.B. Strelets, C.J. Tabone, J. Thurmond, F. Consortium, FlyBase: updates to the Drosophila melanogaster knowledge base, Nucleic Acids Res. 49 (2020) D899–D907. https://doi.org/10.1093/nar/gkaa1026.

